# MESOMELIC DYSPLASIAS ASSOCIATED WITH THE *HOXD* LOCUS ARE CAUSED BY REGULATORY REALLOCATIONS

**DOI:** 10.1101/2021.02.01.429171

**Authors:** Christopher Chase Bolt, Lucille Lopez-Delisle, Bénédicte Mascrez, Denis Duboule

## Abstract

Some human families display severe shortening and bending of the radius and ulna, a condition referred to as mesomelic dysplasia. Many of these families contain chromosomal rearrangements at 2q31, where the human *HOXD* locus maps. In mice, the dominant X-ray-induced *Ulnaless* inversion of the *HoxD* gene cluster produces a similar phenotype suggesting that the same mechanism is responsible for this pathology in humans and mice. Amongst the proposed explanations, the various alterations to the genomic structure of *HOXD* could expose *Hoxd13* to proximal limb enhancers, leading to its deleterious gain-of-expression in the embryonic forelimb. To assess this hypothesis, we used an engineered 1Mb large inversion including the *HoxD* gene cluster, in order to position *Hoxd13* within a chromatin domain rich in proximal limb enhancers. We show that these enhancers contact and activate *Hoxd13* in proximal cells, concomitant to the formation of a mesomelic dysplasia phenotype. A secondary mutation in the coding frame of the HOXD13 protein in-*cis* with the inversion completely rescued the limb alterations, demonstrating that ectopic HOXD13 is indeed the unique cause of this bone anomaly. Single cell expression analysis and evaluation of HOXD13 binding sites in cells from this ectopic expression domain suggests that the phenotype arises primarily by acting through genes normally controlled by HOXD13 in distal limb cells. Altogether, these results provide a conceptual and mechanistic framework to understand and unify the molecular origins of human mesomelic dysplasia associated with 2q31.

## INTRODUCTION

Several human families displaying shortened and bent forearm bones have been reported with large chromosomal rearrangements in the q31 band of chromosome 2, a region containing the *HOXD* gene cluster (Cho et al., 2010; Kantaputra et al., 2010; Le Caignec et al., 2019; Peron et al., 2018). Although they are correlated, the potential involvement of *HOX* genes in causing these limb dysmorphias (mesomelic dysplasias) has never been confirmed, despite various studies in mice indicating that some *Hox* mutations can reproduce this condition. For example, the loss-of-function of either *Hoxd11* or *Hoxa11* individually produced mild phenotypes in the forelimbs (Davis and Capecchi, 1994; Small and Potter, 1993), but when these paralogous null alleles were combined, a severe mesomelic dysplasia of the radius and ulna appeared, resembling human arms with 2q31 alterations (Boulet and Capecchi, 2004; Davis et al., 1995; Wellik and Capecchi, 2003). However, none of the human families evaluated by genomic analyses showed any mutations affecting *HOX* gene bodies, suggesting that the limb malformations were likely to result from mutations interfering with the highly coordinated regulation of *HOXD* gene transcription during early limb development (Andrey et al., 2013; Tarchini and Duboule, 2006).

A strong mesomelic dysplasia (MD) was reported in mice carrying the X-ray-induced mutation *Ulnaless*, an inversion mapping to the murine *HoxD* locus (Davisson and Cattanach, 1990; Spitz et al., 2003). Evaluation of *Hox* transcripts in the limbs of *Ulnaless* mutant embryos revealed the ectopic presence of *Hoxd13* transcripts in the presumptive cellular domain for the radius and ulna, but also gave conflicting results on the down-regulation of both *Hoxa11* or *Hoxd11* (Hérault et al., 1997; Peichel and Vogt, 1997). Because *Hoxd13* is normally transcribed only in the most distal cells of the developing limb buds, where digits are formed (Dollé et al., 1991), the possibility that mesomelic dysplasias in both human and mice are caused by a deleterious *Hoxd13* gain-of-function in the proximal domain, where long bones of the forearm normally develop, was put forward (Hérault et al., 1997). This hypothesis was supported by the dominant nature of these malformations in both the human conditions (Kantaputra et al., 1992) and the mouse *Ulnaless* mutant (Davisson and Cattanach, 1990), the latter being mostly homozygous lethal (Hérault et al., 1997; Peichel and Vogt, 1997).

Extensive chromosome engineering at the murine *HoxD* locus has shed light on the complex regulation of these genes during limb development. The gene cluster is flanked by two ca. 1Mb regulatory landscapes. Centromeric to the cluster (on the side of *Hoxd13*), a range of digit-specific enhancers regulate the transcription of *Hoxd13* to *Hoxd10* in the most distal cells of the growing limb bud (Montavon et al., 2011) (Figure 1a, C-DOM). On the other side of the gene cluster, a series of proximal limb enhancers activate *Hoxd9* to *Hoxd11* in developing forearm cells (Andrey et al., 2013)(Figure 1a, T-DOM). This bimodal type of regulation is organized by the presence of an insulation boundary localized between *Hoxd11* and *Hoxd12* which is established by several bound CTCF proteins (Rodríguez-Carballo et al., 2017). Under normal conditions, this strong insulation boundary prevents the activation of *Hoxd13* in forearm cells by proximal limb enhancers. In the *Ulnaless* allele, the *HoxD* cluster is inverted (Spitz et al., 2003)(see Figure 1a) and as a consequence, *Hoxd13* is brought into the vicinity of known forearm enhancers, putatively explaining its ectopic activation in proximal limb cells. Since the semi-dominant gain of *Hoxd13* expression coincides with a phenotype that mimics the combined loss of both *Hoxd11* and *Hoxa11* in the proximal limb, it was proposed that the presence of the HOXD13 protein would either directly repress the transcription of *Hox11* genes (Peichel and Vogt, 1997), or inhibit the function of group 11 HOX proteins through a dominant negative mechanism referred to as ‘posterior prevalence’ (Duboule and Morata, 1994).

**Figure 1.**
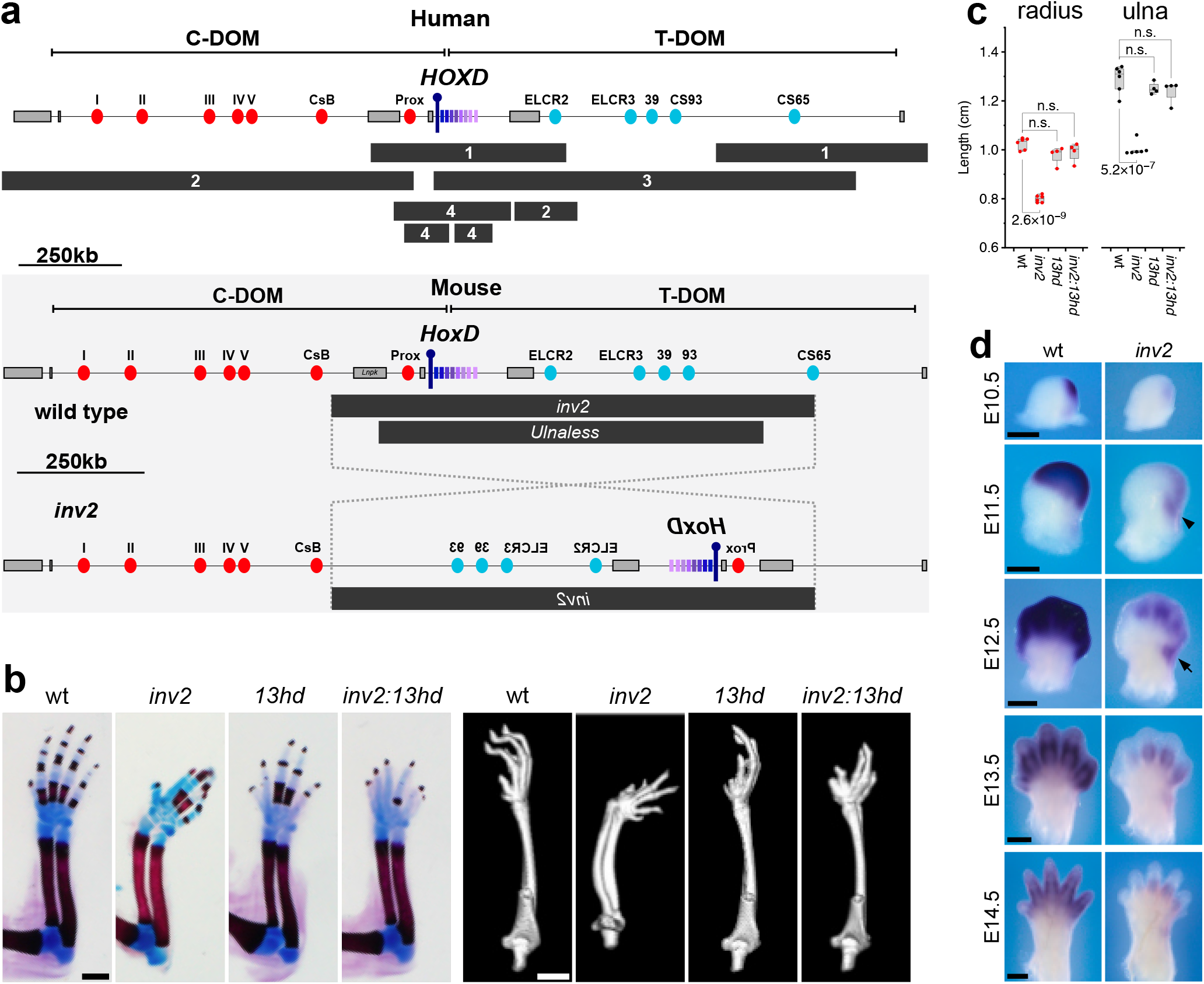
Inversion of the *Hoxd* gene cluster induces limb mesomelic dysplasia. (a) The top panel shows a map of the human *HOXD* locus with the positions of mapped chromosomal rearrangements (black bars) leading to mesomelic dysplasia. The numbers inside the black bars refer to the original references: 1 (Kantaputra et al., 2010), 2 (Peron et al., 2018), 3 (Cho et al., 2010), 4 (Le Caignec et al., 2019). The panels below (grey box) depict the wild type mouse locus (top) and the structure of the *Hoxd^inν2^* inverted allele. In both murine and human loci, the proximal (blue ovals) and distal (red ovals) limb enhancers are indicated, located on either side of the cluster, within the T-DOM and C-DOM landscapes, respectively. *Hoxd* genes are indicated in shades of magenta, with a purple pin to indicate the position of *Hoxd13*. The location of the mouse inversion allele *Ulnaless* is indicated below the *inν2* allele. (b) Newborns skeletal stains (left, scale is 1mm) and adults micro-CT scans (right, scale is 2.5mm) showing mesomelic dysplasia associated with the *inν2* allele. Inactivation of *Hoxd13* in-*cis (inν2:13hd*) completely rescues the alteration. With the exception of the wild types, the *inν2, 13hd*, and *inν2:13hd* mutants all show digit alterations due to significant reduction of *Hoxd13* transcripts in the distal limb (see also Supplementary Figure 1 and Supplemental Video 1). (c) Quantification of bone length based on CT scans showing that *inν2* radius and ulna are significantly shorter than wild type control bones. The bones of *13hd* control mice do not show proximal limb defects. *inν2:13hd* radius and ulna bones are identical in length to control. Box plots are interquartile range, and P-values were determined by Welch’s unequal variances *t*-test, (wt n = 6, *inν2* n = 6, *13hd* n = 4, *inν2:13hd* n = 4), n.s. is not significant. (d) Time course *In situ* hybridizations for *Hoxd13* mRNAs showing both the decrease of transcripts in distal cells and the ectopic proximal expression domain (arrow), most visible at E11.5 and E12.5. Scale bars are 0.5mm.

The need to prevent expression of *Hoxd13* in proximal limb bud cells was further documented by forcing expression in the whole limb bud. Early and strong expression of the transgene completely ablated limb formation proximal to the hands and feet (Goff and Tabin, 1997; Williams et al., 2006; Yokouchi et al., 1995). Another chromosomal rearrangement at the *HoxD* locus showed that a late and weak gain of *Hoxd13* transcription in the proximal limb was enough to shorten the length of the radius and ulna (Tschopp and Duboule, 2011).

While the evidence from these mouse models suggested the gain of HOXD13 as causative, several key questions remained to be answered to turn this hypothesis into an explanation. For instance, how is *Hoxd13* transcription gained in proximal limb cells? Is the gained HOXD13 protein really the cause of the observed alterations and if yes, does ectopic HOXD13 produce these alterations by directly down-regulating *Hox11* transcription or does HOXD13 interfere with HOX11 protein activity in a dominant negative manner? To answer these questions, we used a novel chromosomal inversion in mice similar to the *Ulnaless* rearrangement, yet with slightly different breakpoints leading to a milder gain of *Hoxd13* expression and accompanying phenotype. This strategy allowed us to raise a hypomorph mutant strain and produce homozygous mutant embryos and adults amenable to molecular analyses. By inducing a secondary mutation in-*cis* with the inversion, we demonstrate that the gain of expression of *Hoxd13* is indeed the sole reason for the mesomelic dysplasia phenotype and that it is caused by the abnormal genomic proximity between this gene and native *HoxD* cluster proximal limb enhancers. Furthermore, single cell RNA-seq and protein binding analyses strongly suggest that the effect of ectopic HOXD13 protein is mediated by its binding in proximal cells, to sites that are normally occupied by HOXD13 in distal cells and by HOX11 proteins in proximal cells, together with a partial reduction in the transcription of *Hoxa11*, but not *Hoxd11*. These results allow us to present an inclusive molecular explanation for the reported human 2q31 mesomelic dysplasias.

## RESULTS

### A mouse model for limb mesomelic dysplasia

Human mesomelic dysplasias have been associated with the *HOXD* locus (Fujimoto et al., 1998; Le Caignec et al., 2019; Peron et al., 2018; Spitz et al., 2002; Ventruto et al., 1983)(Figure 1a), but the gene bodies were not affected. Instead, the physical relationship with the flanking regulatory regions was modified, suggesting a potential impact of chromosomal rearrangements upon the long-range regulation of these genes during early limb development (Andrey et al., 2013; Kragesteen et al., 2018; Le Caignec et al., 2019). While the murine *Ulnaless* X-ray-induced inversion is an excellent proxy for these conditions, the severity of its effects and the early homozygous lethality, perhaps due to a breakpoint in the *Lnpk* gene (Figure 1a, *Ulnaless*), made the use of these mice difficult for further analyses and genome editing. We circumvented these problems by using a novel *HoxD* inversion (*HoxD^inν2^* mm10 chr2:74477755-75441001), which was engineered with a 5’ breakpoint within C-DOM, just downstream of the *Lnpk* gene, whereas the 3’ breakpoint was positioned telomeric to the gene cluster within the proximal limb regulatory domain (Figure 1a, *inν2*). As a consequence, the *Lnpk* gene remained intact and most proximal limb enhancers were inverted along with the gene cluster (Figure 1a, bottom panel). Since these latter enhancers were likely responsible for the strong gain of *Hoxd13* expression in the *Ulnaless* inversion, this new inversion was expected to produce a weaker phenotype and viable homozygous specimen, allowing us to carry out the necessary analyses.

This inversion was produced by the STRING approach (Spitz et al., 2005) and mice were born at a Mendelian ratio, without any detectable limb anomaly in the heterozygous condition, unlike the *Ulnaless* allele. However, F2 mice homozygous for the *HoxD^inν2^* (hereafter *inν2*) inversion displayed a clear abnormal morphology of their forelimbs, which was accompanied by a detectable problem in walking. This abnormal phenotype, reminiscent of a mild limb mesomelic dysplasia was fully penetrant. The analysis of skeletal preparations revealed that the radius and ulna were ill-formed, shortened, bent towards the posterior aspect and rotated approximately 90° along their length with respect to the position of the humerus. These combined alterations led to the observed abnormal angle between the hands and the forearm (Figure 1b). This phenotype, only observed in *HoxD^inν2/inν2^* mice, was confirmed and further assessed after micro-CT scans of several mutant and control skeletons (Figure 1b, right panels). CT scans allowed for precise measurements of bone lengths and revealed a significant shortening (ca. 20%, p < 1e-6) of both radius and ulna (Figure 1c). In addition, the digits of *inν2* mice were abnormal, showing a pattern reminiscent of a partial loss of function of *Hoxd13* (Figure 1b, Supplementary Video 1). Therefore, the *HoxD^inν2^* allele produced living homozygous mice with a light yet fully penetrant and significant limb mesomelic dysplasia.

### Ectopic *Hoxd13* transcription and phenotypic rescue through a secondary mutation

We determined whether the *inν2* allele had expectedly induced an ectopic expression of *Hoxd13* in developing forearm cells, as for the *Ulnaless* inversion, by performing time-course analyses of *Hoxd13* mRNAs by whole-mount *in situ* hybridization (WISH)(Figure 1d). During the earliest phase of *Hoxd13* expression, we observed a weak staining when compared with wild type limb buds. Shortly after, by E11.5 when the proximal and distal limb domains begin to separate, a clear *Hoxd13* signal was apparent in the posterior-distal portion of the nascent proximal limb domain of *inν2* mutants and absent from control littermates (Figure 1d black arrowhead).

From E12.5 to E14.5, the ectopic domain of *Hoxd13* mRNA continued to be detected in the posterior-distal part of the proximal limb domain separated from the distal domain by a thin strip of low-expressing cells, i.e. exactly matching the position of the future distal end of the ulna (Figure 1d, arrow). While this ectopic domain was fully penetrant, it was clearly weaker and smaller than in the *Ulnaless* mutant limb buds (Hérault et al., 1997; Peichel and Vogt, 1997), probably due to the fact that the strong proximal limb enhancer CS65 (Andrey et al., 2013) is not adjacent to *Hoxd13* in the inν2 allele, unlike in *Ulnaless*, instead leaving only a few putative enhancer sequences (Andrey et al., 2013) at their initial positions (see below). By E13.5 the transcription of *Hoxd13* was diminished in the proximal and distal limbs of *inν2* limbs.

To demonstrate that this localized ectopic domain of *Hoxd13* mRNAs was indeed causative of the limb mesomelic dysplasia phenotype, we used a CRISPR-Cas9 approach to induce a secondary mutation in-*cis* with the inverted chromosome to functionally inactivate the HOXD13 protein (Supplementary Figure 1a). We induced a 7bp deletion causing a frame shift mutation N-terminal to the nuclear localization signal and homeodomain of HOXD13, the latter domain being necessary for binding to the major groove of target DNA sites. At the same time, the same 7bp deletion was also isolated on the wild type chromosome as a control allele (Supplementary Figure 1b). Mutations disrupting formation of the HOXD13 homeodomain were shown to induce a loss-of-function phenotype in the distal limbs and indeed mice homozygous for this *Hoxd13^hd^* mutation alone displayed the well described *Hoxd13* loss-of-function phenotype in their digits (Dolle et al., 1993)(Figure 1b). While mice homozygous for this *Hoxd13^hd^* mutation in-*cis* with the *HoxD-^inν2/inν2^* inversion (*HoxD^inν2/inν2^:Hoxd13^hd/hd^*) also displayed the expected loss-of-function phenotype in their digits, the mesomelic dysplasia was completely rescued with full penetrance, leading to normal forelimbs as verified by both skeletal staining and micro-CT analyses (Figure 1b and c, Supplementary Video 1). This result demonstrated that the gain of *Hoxd13* function in proximal cells was indeed the unique and proximal cause of limb mesomelic dysplasia.

### Topological reconfiguration of enhancer-promoter interactions after inversion

Expression of *Hoxd* genes during limb development is controlled by two large regulatory landscapes (Figure 1a, C-DOM and T-DOM) which also match two topologically associating domains (TADs)(Dixon et al., 2012; Nora et al., 2012; Sexton et al., 2012) that flank the gene cluster (Figure 2a). The insulation boundary between these two TADs (Rodríguez-Carballo et al., 2017) relies upon the presence of multiple CTCF sites that split the gene cluster into two distinct parts, separating *Hoxd13* and *Hoxd12* from the other genes of the cluster (Bolt and Duboule, 2020). *Hoxd13* is located centromeric to the TAD boundary (Figure 2a, purple pin) and responds exclusively to the various digit enhancers localized in C-DOM (Figure 2a, red ovals), which are active only in distal limb cells. In contrast, enhancers within T-DOM (Figure 2a, blue ovals) are active and promote *Hoxd9, Hoxd10, Hoxd11*, and *Hoxd12* transcription in the proximal limb. In the *inν2* allele, the breakpoints of the 963 kb inversion lie on either side the *HoxD* cluster, positioned within each one of the two TADs (Figure 2a dashed lines). In order to assess the regulatory reallocations induced by these topological modifications, we collected cells from wild type and *inν2* mutant proximal and distal forelimbs and measured DNA-DNA interaction frequencies by Capture Hi-C (CHi-C). The captured sequences were aligned to a reconstructed *inν2* mutant genome (Fig 2a, bottom profile).

**Figure 2.**
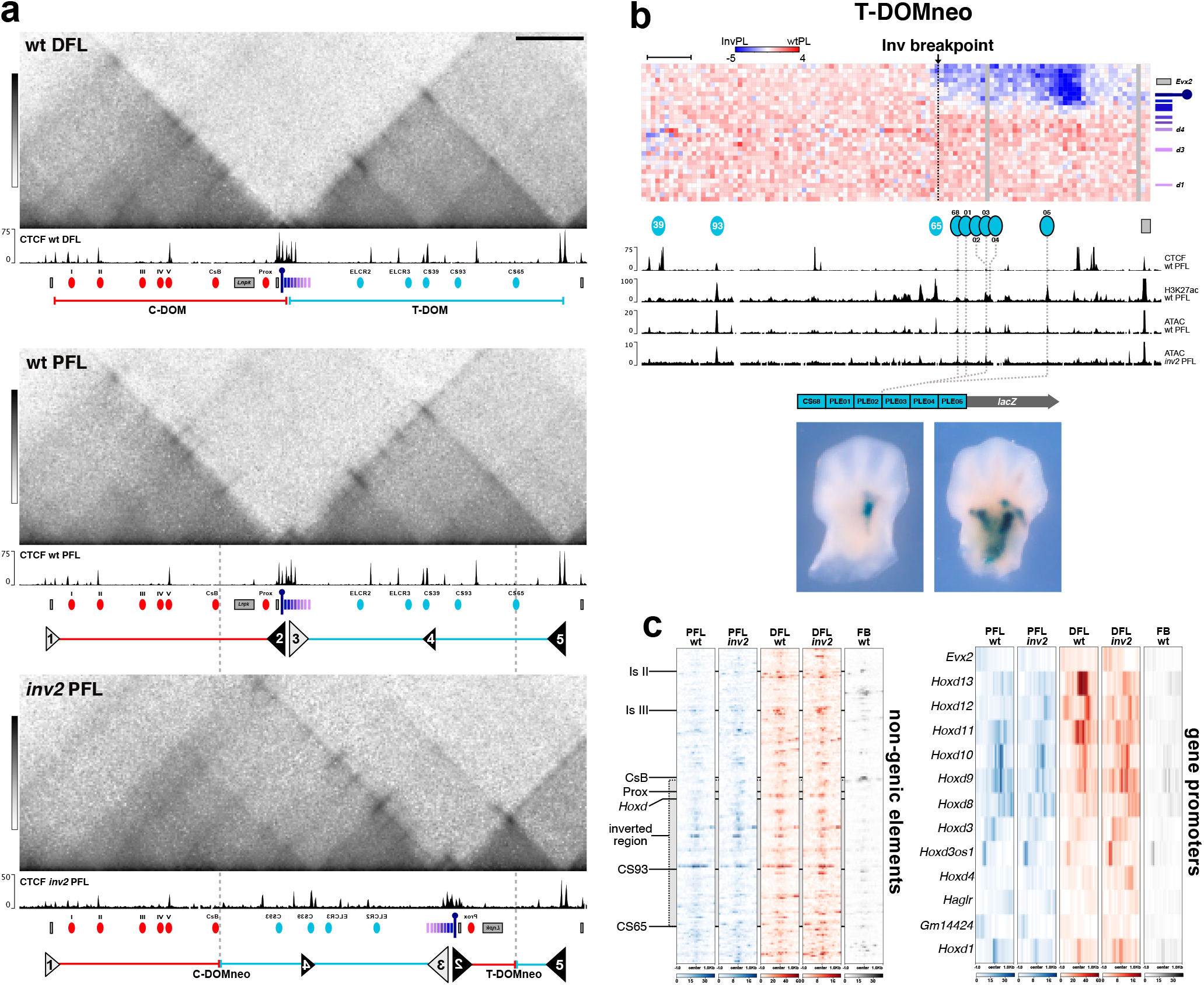
*Hoxd13* establishes new contacts with proximal limb enhancers in the *inν2* allele. (a) Capture Hi-C using E12.5 wild type distal (top) and proximal (center) limb cells, as well as mutant *HoxD^inν2/inν2^* proximal limb cells (bottom). Bin size is 5kb, color scale is log transformed. Wild type samples are mapped to wild type chr2 and *inν2* is mapped on a reconstructed mutant chromosome 2 (mm10). Below each heatmap are CTCF CUT&RUN tracks produced from the indicated tissues and the black and white triangles indicate the orientations of large (big triangles) or reduced (small triangles) groups of CTCF sites. The grey dashed lines between the wildtype PFL and *inν2* PFL indicate the breakpoints of the inversion. In the *inν2* allele, the observed changes in chromatin contacts matches the expectations based on groups of CTCF sites with convergent orientations. The groups are labelled from 1 to 5 to facilitate the reading of the profile after inversion. *Hoxd* genes are colored in shades of magenta and the position of *Hoxd13* is indicated with a purple pin. (b) The top panel shows the subtraction between the CHi-C contacts established by *Hoxd13* and proximal limb enhancers in wild type and in *HoxD^inν2/inν2^* proximal limb cells (each bin is 5kb). Blue bins represent contacts more commonly observed in the *inν2* allele and are concentrated in the T-DOMneo domain, starting right after the position of the breakpoint (vertical line). The mapping is shown on the wildtype chromosome for clarity. The tracks below show CTCF CUT&RUN, H3K27Ac ChIP (Rodríguez-Carballo et al., 2017) and ATAC using E12.5 wild type or *inν2* proximal forelimb cells, mapped onto wildtype mm10. Previously characterized proximal limb enhancers are indicated by blue ovals below the heatmap. Putative proximal limb enhancers that gain contact with *Hoxd13* in the *inν2* allele are indicated by blue ovals with a black border. These elements (CS68 and PLE01 to 05) were identified through the H3K27Ac and ATAC profiles in wild type and *inν2* proximal forelimbs and were cloned into a single *lacZ* reporter transgenic construct. Representative staining patterns in forelimb buds are shown below (see Supplementary Figure 2e). All embryos that stained (7 of 7) produced a variation of the proximal limb staining. (c) Comparison of ATAC-seq datasets between the profiles in control and *inν2* proximal forelimbs. The heatmaps on the left are peaks from ATAC samples mapped onto non-genic elements at the *HoxD* locus, illustrating the high similarity between samples of the same tissue, regardless of genotype. The regions corresponding to the gene bodies of *Lnpk*, *Mtx2* and the region from *Evx2* to *Hoxd1* have been removed. The heatmaps on the right are the ATAC profiles over accessible gene promoters in the *HoxD* cluster. The PFL samples are generally similar while the DFL samples show more difference, in particular on *Hoxd13* and *Hoxd11* promoters.

In distal limb cells of E12.5 control embryos, the chromatin conformation displayed the well-characterized topology of the *HoxD* locus containing two TADs separated by the CTCF-dependent insulation boundary present within the gene cluster (Rodríguez-Carballo et al., 2017). In the *inν2* allele, a major redistribution of contacts was observed, which could be explained by the reorganization of the various CTCF insulation boundaries present over the entire locus. We defined insulation boundaries as a concentration of occupied CTCF sites capable of producing a bidirectional boundary effect in normal limb cells. Five such boundaries were identified, schematized as triangles and labelled from 1 to 5. The orientation of the sites is reflected by the orientation and color of the triangles, where sizes indicate the strength of the boundary effect (Figure 2a, middle panel).

On the inversion allele, the *HoxD* insulation boundary (Figure 2a, triangles 2 and 3) was inverted and moved closer to the telomeric end of T-DOM. From this new position, one boundary element (triangle 2) established contacts with the existing telomeric boundary (triangle 5) to induce formation of a new TAD (T-DOMneo). This small and dense domain contained *Hoxd13*, a single digit enhancer (Prox)(Gonzalez et al., 2007), and a few putative proximal limb enhancer elements that remain in place since they are located beyond the telomeric inversion breakpoint (the blue portion of T-DOMneo). All other proximal limb enhancers were relocated to the other side of the locus (Figure 2a, between boundary triangles 1 and 3) where they also formed a new TAD structure with the large portion of C-DOM that had stayed at its initial position (Figure 2a, C-DOMneo between boundaries 1 and 3). Thus, the novel C-DOMneo regulatory landscape contained the majority of functional proximal and distal limb enhancers (Supplementary Figure 2b).

To determine if the reconfiguration of the regulatory landscapes had altered the expression of other *Hoxd* genes, we evaluated their expression by WISH. In *inν2* mutants we found a minor increase in *Hoxd11* and *Hoxd12* expression in the proximal limb buds (Supplementary Figure 2c) likely resulting from the same change in contacts affecting *Hoxd13* expression. In contrast, we observed a reduction of *Hoxd12, Hoxd11*, and *Hoxd10* in the distal limb compartment, certainly resulting from the increased distance between these genes and their distal limb enhancers within the C-DOMneo configuration. The fact that only the Prox distal limb enhancer remained with *Hoxd13* after inversion explained the severely reduced transcription of *Hoxd13* in distal limb cells and the associated phenotype.

We then looked at T-DOMneo to determine if *Hoxd13* expression in proximal cells after inversion could be linked to increased contacts with the portion of the proximal limb regulatory landscape that had not been inverted (Figure 2b). In the inverted allele, *Hoxd13* indeed established significant interactions with a region that was labelled by H3K27 acetylation in proximal limb cells at this stage (Beccari et al., 2016), and which displayed several accessible chromatin regions, as assayed by ATAC-seq (Figure 2b)(Buenrostro et al., 2013). The presence of putative regulatory elements within the T-DOMneo suggested that these elements may be responsible for the ectopic expression of *Hoxd13* in the PL.

To confirm that this particular region carried some proximal regulatory activity, we identified six regions showing H3K27ac signal in wild type proximal limbs, ATAC peaks in the *inν2* samples, and no apparent CTCF binding. These six putative enhancer elements, CS68 and Proximal Limb Enhancers PLE01 to PLE05 (Figure 2b and Supplementary Table 1) were generated as a concatenated element and positioned 5’ to a *lacZ* reporter construct. All transgenic embryos showing β-gal staining were positive in the proximal limb (7 out of 7), however the size of the expression domain varied from a small discrete patch to several larger regions extending towards the distal boundary of the proximal limb domain (Figure 2b and Supplementary Figure 2e). While the domains were not always completely overlapping with the ectopic *Hoxd13* domain, these results confirmed that DNA sequences at the vicinity of *Hoxd13* in the *inν2* configuration are active in proximal limb cells, where they also significantly increased their contacts with *Hoxd13*. Noteworthy, other sequences located telomeric to TAD boundary #5 were shown to contribute to *Hoxd* gene expression in a posterior patch of proximal cells (Yakushiji-Kaminatsui et al., 2018), which together with the sequences reported here, may all contribute to the ectopic expression of *Hoxd13* in the inverted allele.

We then evaluated if the structural changes observed in the CHi-C may have contributed to the ectopic expression of *Hoxd13* in proximal limb cells by altering the enhancer status of the two regulatory landscapes. To do so, we used our ATAC-seq datasets to identify any changes in chromatin accessibility in proximal and distal limbs of both genotypes and a wild type forebrain as control (Fernandez-Guerrero et al., 2020). First, we selected the ATAC-seq peaks corresponding to non-genic elements located throughout the locus outside of the gene cluster. We then evaluated them visually using heatmaps and found a high correspondence between the accessibility status and the tissue of origin (Figure 2c left). This was confirmed by hierarchical clustering analysis of the non-genic elements (Supplementary Figure 2d left panel). However, when we evaluated the accessible gene promoters in the HoxD cluster, there were only minor differences between the PFL samples, but noticeable differences between the DFL samples, especially on *Hoxd13* and *Hoxd11* (Figure 2c right). We repeated the clustering analysis on these regions and observed that they no longer clustered by tissue of origin, rather both *inν2* samples clustered closely with the wild type PFL sample (Supplementary Figure 2d), and the differences are most apparent in the genomic coverage map of the gene cluster (Supplementary Figure 2a ATAC). These changes in promoter accessibility, in particular *Hoxd11* and *Hoxd13* in the DFL, mirror the reduction in expression of these genes observed by WISH, and support the conclusion that the inversion did not change the activity status of the regulatory landscapes. Instead, the inversion altered the relationship between the genes and their enhancers by forming a new three-dimensional structure that restricted 5’ genes from their normal C-DOM enhancers, and simultaneously introduced them to proximal limb enhancers, which they are normally insulated from.

### Excluding *Hox11* transcriptional down-regulation as the secondary cause of mesomelic dysplasia

Having established the mechanism leading to the gain of expression of the *Hoxd13* gene in proximal limb cells and the fact that ectopic HOXD13 in these cells is the sole cause for the limb deformities, we addressed the potential mechanisms through which the protein may achieve its deleterious effect. The limb alterations produced by either the *Ulnaless* or the *inν2* alleles are both similar to the limb phenotypes found in mice with significant reductions in *Hoxa11* and *Hoxd11* transcription (remaining expression <50% in *Hoxa11* and <50% in *Hoxd11*)(Davis et al., 1995). One proposed explanation is that ectopic HOXD13 may abrogate transcription of *Hoxa11* and *Hoxd11* in proximal limb cells so that the mesomelic dysplasia phenotype converges towards a combined *Hox11* loss-of-function phenotype (Peichel and Vogt, 1997). However, another study did not observe any substantial change in *Hox11* transcription in proximal *Ulnaless* limbs (Hérault et al., 1997).

We revisited these results by performing *in situ* hybridizations for *Hoxa11* and *Hoxd11* but imaged the embryos in a time course through the linear phase of color development when differences in staining are more apparent (Figure 3a and Supplementary Figure 3a). At E12.5, we observed that *Hoxa11* transcripts are slightly but clearly reduced in a small region of the proximal limb of *inν2* mutants corresponding to the ectopic *Hoxd13* domain (arrow in Figure 3a and arrowhead in Supplementary Figure 3a). Under the same conditions, we also observed a similarly small but consistent increase of *Hoxd11* transcripts at the same position (arrowhead in Figure 3a and Supplementary Figure 3a). Even so, this partial reduction of *Hoxa11* transcripts by *Hoxd13* is not sufficient to explain the observed phenotype, especially without an at least equivalent or greater reduction of *Hoxd11* transcripts.

**Figure 3.**
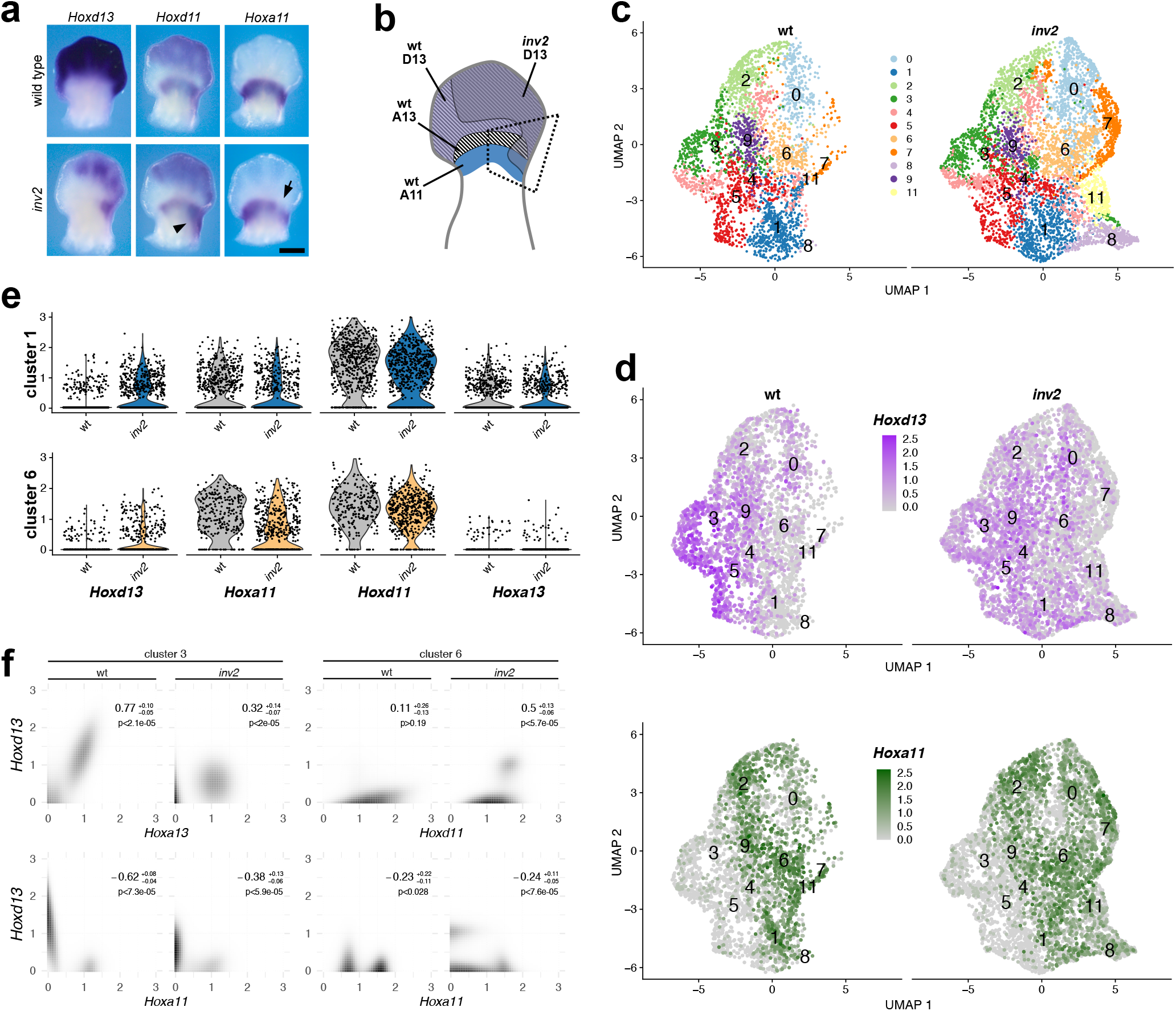
Single cell RNA-seq analysis of the *inν2* limb ectopic *Hoxd13* domain. (a) *In situ* hybridizations comparing the ectopic gain of *Hoxd13* in the proximal forelimb to the expression of *Hoxa11* and *Hoxd11*. A small decrease in *Hoxa11* transcripts was scored in the posterior portion of proximal limb domain (arrow), whereas *Hoxd11* staining is slightly increased in the posterior and distal portion of the proximal forelimb (arrowhead) when compared with wild type limb buds. Scale bar is 0.5mm (see also Supplementary Figure 3a). (b) Scheme of an E12.5 limb with the regions of gene expression for wild type (*Hoxa11* in blue, *Hoxd13* in purple and *Hoxal3* in black and white stripes, as well as the approximate location of *Hoxd13* mRNAs in the *inν2* allele (grey). The region outlined by the black dashed quadrangle was dissected for single cell RNA-seq analysis. (c) UMAP representation of the primary cellular clusters as determined by Seurat. All clusters were found in both genotypes, but clusters 7, 8, and 11 were present in greater proportion in the *inν2* samples than in the wild type. (d) The top panel is the UMAP representation for the main cluster with cells expressing *Hoxd13* indicated in shades of purple according to the level of expression, and *Hoxa11* in shades of green. In the wild type sample, *Hoxd13* expression was primarily limited to the distal limb clusters (3, 4, 5, and 9). With the *inν2* sample, however, the expression was reduced in distal limb clusters while it increased in the proximal limb clusters 1 and 6. (e) Clusters 1 and 6 were the only ones that expressed both *Hoxa11* and *Hoxd11*, along with a gain of *Hoxd13* in the *inν2* configuration. The violin plots represent the detected transcript expression for each cell in that cluster. (f) Heatmaps representing the fitted distribution of cells for the different levels of expression of *Hoxd13* on the y-axis and *Hoxa11*, *Hoxd11* or *Hoxal3* on the x-axis. The color scale is log transformed in order to see a greater range of frequency. Bins with black represent a high proportion of cells whereas bins with white indicate an absence of cells. In the right corners are indicated the confidence interval of the correlation as well as an estimation of the one-sided p-value (probability that the correlation has the opposite sign).

Because *in situ* hybridizations have a poor cellular resolution and are difficult to quantify, we implemented single-cell RNA-seq to evaluate a potential correlation between ectopic HOXD13 and a reduction in the amount of *Hox11* transcripts. We micro-dissected a region including the ectopic patch of *Hoxd13* mRNAs in both *inν2* and wild type limbs (Figure 3b, dashed quadrangle) and processed them for single cell RNA-seq using the 10X Chromium platform with 3.1 chemistry. We sequenced 5006 cells from one wild type and two *inν2* biological replicates producing 4535 and 4315 cells, with a mean number of reads per cell between 60’000 and 80’000, and analyzed with the Seurat package (Butler et al., 2018). Clustering analysis displayed in a 2-dimensional UMAP identified one main group consisting of 11 clusters (Figure 3c). All clusters were identified in both genotypes, but three clusters (7, 8, 11) were present in greater proportions in the *inν2* samples, suggesting no major changes to clustering identities between the two genotypes.

To separate proximal from distal cell clusters, we identified the clusters where *Hoxd13* was strongly expressed in wild type cells (clusters 3, 4, 5, 9), and did the same for *Hoxa11* (clusters 0, 1, 2, 4, 6, 7, 8, 9, 11), which was strongly associated with cells also expressing the proximal limb marker *Shox2* (Figure 3d, Supplementary Figure 3b). We then tried to visually identify clusters which 1) had an increase of *Hoxd13* in the *inν2* configuration, 2) express *Hoxa11* in the wild type, and 3) express other proximal limb markers. We identified two clusters meeting these criteria (clusters 1 and 6)(Figure 3e and Supplementary Figure 3b). In the UMAP, these two clusters reside along the boundary between cells that are distinguished by proximal and distal limb marker genes, yet more closely associate with cells displaying a proximal limb identity (Figure 3c-d and Supplementary Figure 3b-c).

In single cell RNA seq experiments, the proportion of zeros is highly anti-correlated with the mean expression value of the gene (Tung et al., 2017). Accordingly, we used the proportion of zeros as a proxy for the mean expression values to evaluate if the ectopic expression of *Hoxd13* in the proximal limb clusters produced a correlational effect on the expression of *Hoxa11* and *Hoxd11*. We predicted that the expression level of *Hoxd13* is higher in cells where *Hoxd13* is detected, and *vice-versa* for cells where *Hoxd13* is not detected. When we analyzed cells with detectable *Hoxd13* we observed that the proportion of cells with *Hoxa11* was always significantly decreased compared to those cells without *Hoxd13* (Supplementary Figure 3d). The exception to this is cluster 6 in the wild type sample because there are very few cells where *Hoxd13* is detected. Following the same reasoning, we found a strong positive correlation between *Hoxd11* and *Hoxd13* in clusters 1 and 6 of the *inν2* sample.

However, this method can be biased when the number of UMIs is different between samples. In order to remove this bias, we used a new method based on a Markov chain Monte Carlo, to evaluate the inferred true distribution of expression for each gene independently, by cluster and by genotype (see Methods). This new method can be applied to any single gene, determining the confidence interval on the distribution of expression, and a confidence interval on the fold-change between two conditions (Supplementary Figure 3e). Using this approach, we found a two-to-three-fold increase in *Hoxd13* in clusters 1 and 6, and observed a 40 to 55% decrease in *Hoxa11* transcripts compared to wild type cells in the same clusters, and a decrease of 15 to 30% for *Hoxd11*.

This method can be extended to evaluate the distribution of expression for two genes at the same time. First, we evaluated the distal limb control cluster 3 and found a clear anti-correlational between *Hoxd13* and *Hoxa11* (Figure 3f, bottom left panel), matching the previous observation that *Hoxd13* represses the transcription of *Hoxa11* in the distal limb (Beccari et al., 2016; Kherdjemil et al., 2016). In the same cluster we found a positive correlation between *Hoxd13* and *Hoxa13*, which is also not surprising due to the high frequency of these genes being expressed in the same distal limb cells (Fabre et al., 2018). Finally, we evaluated the proximal limb cluster 6 and found a clear anti-correlation between *Hoxd13* and *Hoxa11*. Together, these results support the conclusion that *Hoxd13* reduces the transcription of *Hoxa11* when expressed ectopically in proximal limb cells, yet to an amount that cannot account for the mesomelic dysplasia phenotype. The absence of a similar effect upon *Hoxd11* in the *inν2* was not unexpected since *Hoxd13* and *Hoxd11* transcripts are often present in the same distal cells (Fabre et al., 2018).

### Ectopic HOXD13 binding pattern in proximal limb cells

In proximal limb cells, inappropriate expression of HOXD13 alters the normal expression of *Hoxa11* so it could also influence the expression of other genes important to normal limb formation. It may do this by binding to the same set of DNA sequences that the protein normally binds in distal cells (Desanlis et al., 2020), thus implementing a ‘distal’ program in these proximal cells. Alternatively, HOXA11 and HOXD13 share very similar binding motifs and binding positions, yet their expression is normally within discrete portions of the limb (Desanlis et al., 2020; Jerković et al., 2017). The co-expression of both factors in the same cells may redirect ectopic HOXD13 binding towards positions already bound by HOXA11. To discriminate between these possibilities, we analyzed HOXD13 binding in *inν2* mutant proximal limb cells by CUT&RUN (Skene et al., 2018). The posterior-proximal forelimb (P-PFL) region containing the *Hoxd13* ectopic domain was micro-dissected in duplicate (Figure 4a, dashed triangle). The remaining portion of the limbs were processed for *Hoxd13 in situ* hybridization as controls for the dissection (Supplementary Figure 4a). To determine if HOXD13 binding is altered in the P-PFL, we also generated HOXD13 CUT&RUN from wild type distal forelimb cells and compared these with a previously reported HOXA11 whole forelimb dataset (Desanlis et al., 2020).

**Figure 4.**
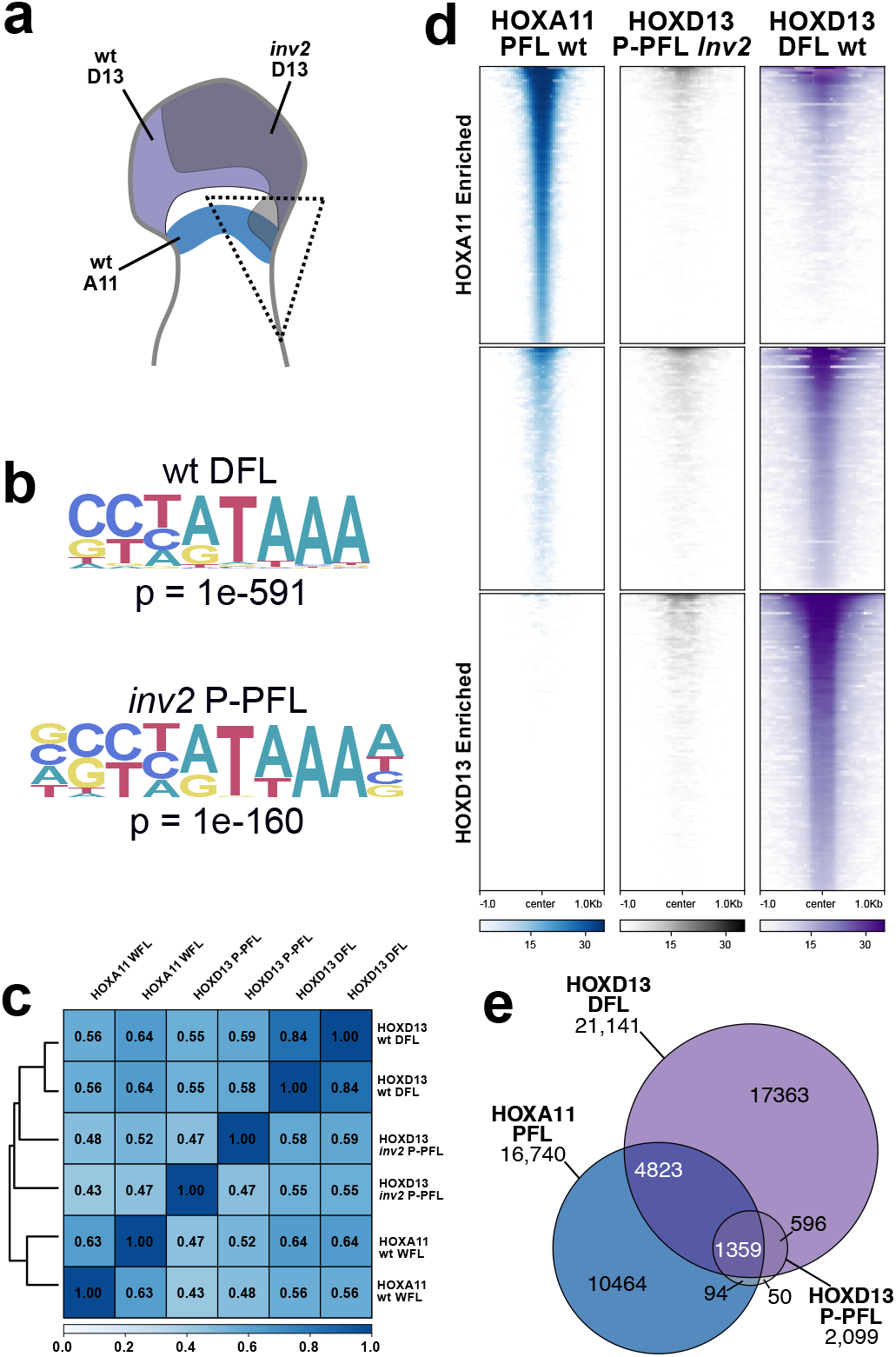
Ectopic HOXD13 binds only to positions bound by HOXD13 in the distal limb. (a) Scheme as in Figure 3, illustrating the dissection for this dataset. (b) HOXD13 consensus binding motif found in E12.5 wild type distal forelimbs (DFL) and *inν2* posterior proximal forelimbs (P-PFL) demonstrates that the protein binds to the same sequences in ectopic domain as it does in the normal distal limb domain, although in the *inν2* samples the motif was extended by one nucleotide on both sides. (c) Pearson correlation clustering on coverage from all replicates used for binding analysis demonstrating that the *inν2* P-PFL HOXD13 coverage most closely cluster with the wild type DFL HOXD13 dataset rather than the coverage of HOXA11 in the PFL. (d) Heatmaps of genomic regions differentially bound by HOXA11 (Desanlis et al., 2020) at E11.5 in the proximal forelimb and HOXD13 at E12.5 in the distal forelimb of wild type embryos. Many of sites preferentially bound by HOXA11 are also bound by HOXD13 in the distal limb, but HOXD13 can also bind to many sites that HOXA11 does not bind to, as previously reported. The HOXD13 binding regions identified in the P-PFL tissue of *inν2* limbs were mapped onto these differentially bound regions. HOXD13 in the P-PFL binds to a portion of sites bound in the wild type distal limb. (e) Euler diagram of binding sites identified as significant peaks in both replicates of each experiment. The majority of HOX13 binding sites in the P-PFL overlap with HOXA11 sites.

We first determined whether the HOXD13 consensus binding motif was the same between the control DFL and the *inν2* P-PFL samples. *De novo* motif finding identified the previously reported motif (Heinz et al., 2010; Jerković et al., 2017; Sheth et al., 2016) in both control and *inν2* samples. In both samples, the HOXD13 motif was the only HOX motif identified among the top five high-scoring results, indicating that proximal forelimb chromatin environment did not alter its preferred binding sequence (Figure 4b). We then performed a hierarchical clustering analysis of HOXD13 binding profiles found in the P-PFL sample to determine if they better match the HOXA11 or the HOXD13 profiles. We found that these samples cluster more closely with the wild type DFL binding profiles than with the HOXA11 PFL samples (Figure 4c). This was confirmed by differential binding analyses (Ross-Innes et al., 2012), where we determined all of the peaks bound preferentially by HOXD13 in control distal cells or by HOXA11 in proximal cells, or not bound preferentially at all, and compared with this the *inν2* P-PFL HOXD 13 peaks. We observed a pattern of HOXD 13 in P-PFL samples that most closely matched the control DFL HOXD13 binding profile although with a much lower signal (Figure 4d). Therefore, it appears that the set of HOXD13 binding sites identified in proximal cells expressing *Hoxd13*, was closely related to the set of binding sites normally occupied by HOXD13 in distal cells, suggesting that the proximal limb cells may have undergone a partial transition to a distal limb identity.

Next, we looked at the percentage of those sites bound by HOXD13 either in control distal or in P-PFL cells, which would also be occupied by HOXA11 in proximal cells (Desanlis et al., 2020). We compared the 24’141 HOXD13 peaks identified in the wild type DFL to the 16’740 HOXA11 binding sites identified in control proximal limbs and found that 6’182 peaks (25% of HOXD13, 37% of HOXA11 peaks) overlapped between the two (Figure 4e). We then evaluated the overlap between the HOXD13 binding sites in proximal cells and the distal forelimb. We found 1’955 sites in common, but the proportion of those that overlap with HOXA11 had increased from 25% to 70% (1’359). This change in proportion suggests that most of the sites uniquely bound by HOXD13 in the distal limb cannot be bound in the proximal limb, either because of the absence of essential co-factors, or because the small pool of HOXD13 factors was preoccupied at sites where HOXA11 is normally bound. Even though the scarcity of starting material may have introduced a technical bias, as illustrated by the total amount of detected peaks, this shift in proportions indicates a change in the binding repertoire towards a condition intermediate to interfering with HOXA11 binding sites and deployment of a distal limb program in the proximal limb cells. This is corroborated by the scRNA-seq experiment where the cells of proximal limb did not show a complete transition to distal limb cell identity.

## DISCUSSION

### *Hoxd13* as the cause of mesomelic dysplasia in mice

In this study, we overcame the difficulty to use the mouse *Ulnaless* allele as a model system to understand the molecular etiology of mesomelic dysplasia (MD), by analyzing a similar yet less drastic condition where slightly different inversion breakpoints generated an ectopic gain of expression of *Hoxd13* in proximal cells, which was weaker and spatially more restricted than in the *Ulnaless* inversion. This weaker expression was due to the removal of most proximal limb enhancers (Figure 5) leaving in place only a few weak proximal limb regulatory sequences. This group of proximal cells expressing *Hoxd13* was nevertheless large enough to induce a fully penetrant MD phenotype in homozygous mice, which otherwise could breed and were thus available for analysis. We took advantage of this to generate a secondary mutation in-*cis* with the inversion, whereby a full loss-of-function of *Hoxd13* was induced. In these mice, the shortening and bending of bones associated with the primary mutation was fully rescued, unequivocally demonstrating the central role of the gained *Hoxd13* in disrupting limb development.

**Figure 5.**
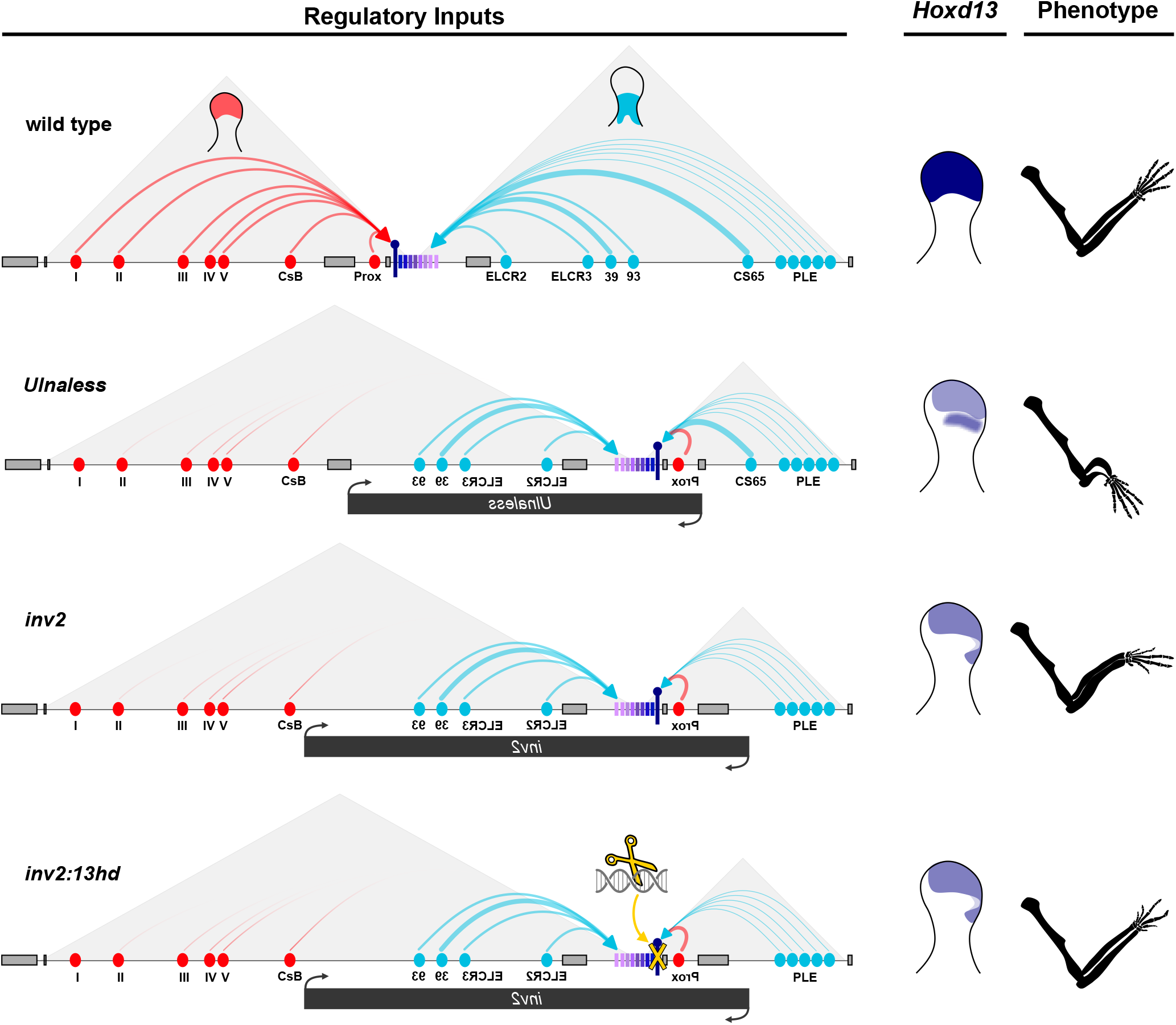
Schematic illustrations of the regulations at work in the various mouse alleles and their effect on *Hoxd13* expression. On top is the control landscape, with distal limb enhancers (red) activating *Hoxd13* (purple pin) in distal cells. On the other side of the cluster, proximal limb enhancers (blue) regulate more telomeric *Hoxd* genes in proximal limb cells. As a result, *Hoxd13* is expressed in distal cells only (right), leading to the wild type phenotype. In the *Ulnaless* inversion, *Hoxd13* is positioned close to both a strong proximal enhancer (CS65) and a series of weaker enhancers (PLE). As a result, it becomes strongly expressed in proximal limb cells, while reduced in distal cells since it is separated from all distal limb enhancers except one (Prox), leading to a light phenotype in digits and strong mesomelic dysplasia. In the *inν2* inversion, *Hoxd13* is not adjacent to CS65 as in *Ulnaless*, but is now under the control of the weaker enhancer series (PLE). The gain of expression in proximal limb cells is weaker and accordingly, the mesomelic dysplasia is not as severe as in *Ulnaless* mice, while the digit phenotype is expectedly comparable. In *inν2:13hd* limb cells (bottom), the transcription of *Hoxd13^hd^* is the same as in *inν2*, yet the mesomelic dysplasia phenotype is completely rescued and the forearms are like wild type. However, because *Hoxd13* is now fully inactivated in distal cells, the digit phenotype is stronger and equivalent to a full *Hoxd13* knock-out (Dollé 1993).

Because the loss of function of both *Hoxa11* and *Hoxd11* induced severe mesomelic dysplasia in mice, it was argued using both murine and human conditions, that various breakpoints around the *HOXD* cluster would lead to the down regulation of the latter two genes thus inducing bone anomalies (Peichel and Vogt, 1997; Peron et al., 2018). Alternatively, it was proposed that while these two genes would remain transcribed, the ectopic presence of HOXD13 protein would functionally interfere with HOXD11 and HOXA11 proteins through a dominant negative effect referred to as ‘posterior prevalence’ (Duboule and Morata, 1994; Hérault et al., 1997). While our datasets produced from both single-cell RNA sequencing and DNA binding analyses of various HOX proteins do not allow us to completely discriminate between these two possibilities, it is clear that the former explanation alone cannot account for the observed MD phenotypes. Indeed, studies involving combined *Hoxa11/Hoxd11* mutations in mice observed mesomelic dysplasia phenotypes when at least half the total dose was removed (Davis et al., 1995). In our single cell RNA-seq experiment we observe in our *inν2* mice a 40 to 55 % reduction of *Hoxa11* and a modest 15 to 30% reduction of *Hoxd11*, which together are not sufficient to elicit the described phenotype. In addition, the weak reduction in *Hoxd11* transcript was not scored in WISH where, if anything, a slight gain was observed.

Therefore, the partial reduction of *Hoxa11* transcription must be potentiated by another effect of ectopic HOXD13. Our differential binding analysis revealed that ectopic HOXD13 was bound to a set of sites that most closely matched those bound by HOXD13 in control distal cells, with more than 25% of the binding sites identified in the proximal limb being sites normally restricted to the distal limb. This suggests that proximal cells may have acquired a more distal identity leading to a reduction in bone length. The recent observation that HOX13 proteins have a pioneer effect (Amândio et al., 2020; Bulajić et al., 2020; Desanlis et al., 2020) provides a potential mechanism for this to occur. On the other hand, in proximal cells ectopically expressing HOXD13, 70% of those sites are normally bound by HOXA11, which suggests that a large part of the change in proximal cells may result from interactions between HOXD13 and HOX11 proteins at these binding sites, potentially through the dominant negative effect of posterior prevalence. This may also apply to other circumstances where the gain of HOXD13 protein led to alterations identical to *Hox11* genes loss of function, for example during the development of metanephric kidneys (Darbellay et al., 2019; Davis et al., 1995; Hoeven et al., 1996).

### An inclusive model for human mesomelic dysplasias at 2q31

Thus far, none of the human mesomelic dysplasias mapping to 2q31 could be directly associated with mutations in *HOXD* genes, but breakpoints were identified and located in the vicinity of the *HOXD* cluster. As a consequence, the various reported deletions, inversions or duplications were generally interpreted as inducing a regulatory loss-of-function of *HOXD* genes, in particular of *HOXD11* (Kantaputra et al., 2010; Peron et al., 2018). Because the regulation of *Hoxd* genes during limb development has been shown to be evolutionarily conserved in amniotes, we can revise previous explanations and propose an inclusive model to account for various mesomelic dysplasia mapped at 2q31, solely based on the abnormal gain of *HOXD13* transcription in proximal limb cells (Supplementary Figure 5).

For instance, the duplication of the *HOXD* cluster reported in Kantaputra et al. (2010) includes the ELCR2 proximal limb enhancer (Rodríguez-Carballo et al., 2020). In the mutant chromosome, a copy of *HOXD13* is in close proximity with this enhancer, regardless of the orientation of the duplicated DNA sequence and will thus receive proximal regulatory inputs (Supplementary Figure 5b, blue arrows), *a fortiori* since one of the duplicated copies of *HOXD13* is now located far from its digits-specific enhancers (red arrows), a situation which was shown in mice to de-sequester *Hoxd13* and re-allocate it towards proximal enhancers on the other side of the cluster (Tschopp and Duboule, 2011). In the case where the duplicated segment would also be inverted, *HOXD13* would even be in closer contact with multiple proximal limb enhancers (Supplementary Figure 5b, top). A similar reasoning applies for the two families described in Le Caignec et al. (2019). In family 1, the duplicated and inverted copy of *HOXD13* is now in close contact with proximal enhancers (Supplementary Figure 5c, top). Likewise, the large inverted duplication mapped in family 2 brings *HOXD13* even closer to proximal limb enhancers (Supplementary Figure 5b, bottom).

Cho et al. (2010) reported a family carrying a duplication extending across the *HOXD* cluster and the proximal limb regulatory landscape, without determining the orientation of the duplicated DNA segment (Supplementary Figure 5d). In both orientations, however, the result is that the duplicated copy of *HOXD13* is de-sequestered from the distal regulatory landscape (red arrows) and is now licensed to interact with proximal enhancers (blue arrows). The same mechanism likely underlies the case reported by Peron et al. (2018), which involves two deletions either in *cis* or in *trans*. If in *trans*, the deletion of all distal limb enhancers (Supplementary Figure 5e, deletion 1) would again allow *HOXD13* to interact with proximal limb enhancers (Supplementary Figure 5, top). An almost identical configuration to this had been previously shown in mice where the distal limb enhancers were removed through a large inversion (Supplementary Figure 5e, bottom)(Tschopp and Duboule, 2011). If both deletions had occurred in *cis*, it is possible that the *HOXD* cluster, located in between, was inverted as well (Supplementary Figure 5, middle, see Kragesteen et al., 2018). In this case, *HOXD13* would be positioned in the vicinity of proximal limb enhancers, leading to a strong gain of expression.

This explanatory framework can thus be applied to all reported cases of human mesomelic dysplasia associated with 2q31, so far without any exception. The various positions of *HOXD13* with respect to the proximal limb enhancers as observed in the different causative chromosomal rearrangements, likely explains the variability in the strength of the alterations scored in the human forearms.

## ETHICS APPROVAL

All experiments involving animals were performed in agreement with the Swiss Law on Animal Protection (LPA), under license No. GE 81/14 (to DD).

## DATA AVAILABILITY

All raw and processed datasets are available in the Gene Expression Omnibus (GEO) repository under accession number GSE165495. All scripts necessary to reproduce figures from raw data are available at https://github.com/lldelisle/scriptsForBoltEtAl2021.

## AUTHORS CONTRIBUTIONS

C. C.B.: Designed and conducted experiments, analyzed datasets, formalized results and wrote the paper.

B.M.: Designed, produced, genotyped and helped analyze mouse mutants.

L.D.: Analyzed and evaluated the statistical significance of datasets. Wrote the paper.

D. D.: Designed experiments, transported mice, dissected some limb buds and wrote the paper.

## ACKNOWLEDGEMENTS

We thank Dr. Leonardo Beccari for early insights on this project, Dr. Josef Zakany for assistance with skeletal preparations and analysis, Dr Alexandre Mayran for advice on single cell RNA-seq as well as members of the Duboule laboratories for comments and discussion. We are grateful to Sandra Gitto and Thi Hanh Nguyen Huynh for their help with mice breeding, as well as to Bastien Mangeat and the EPFL Gene Expression Facility staff.

## LEGENDS TO SUPPLEMENTARY FIGURES

**Supplementary Figure 1.**
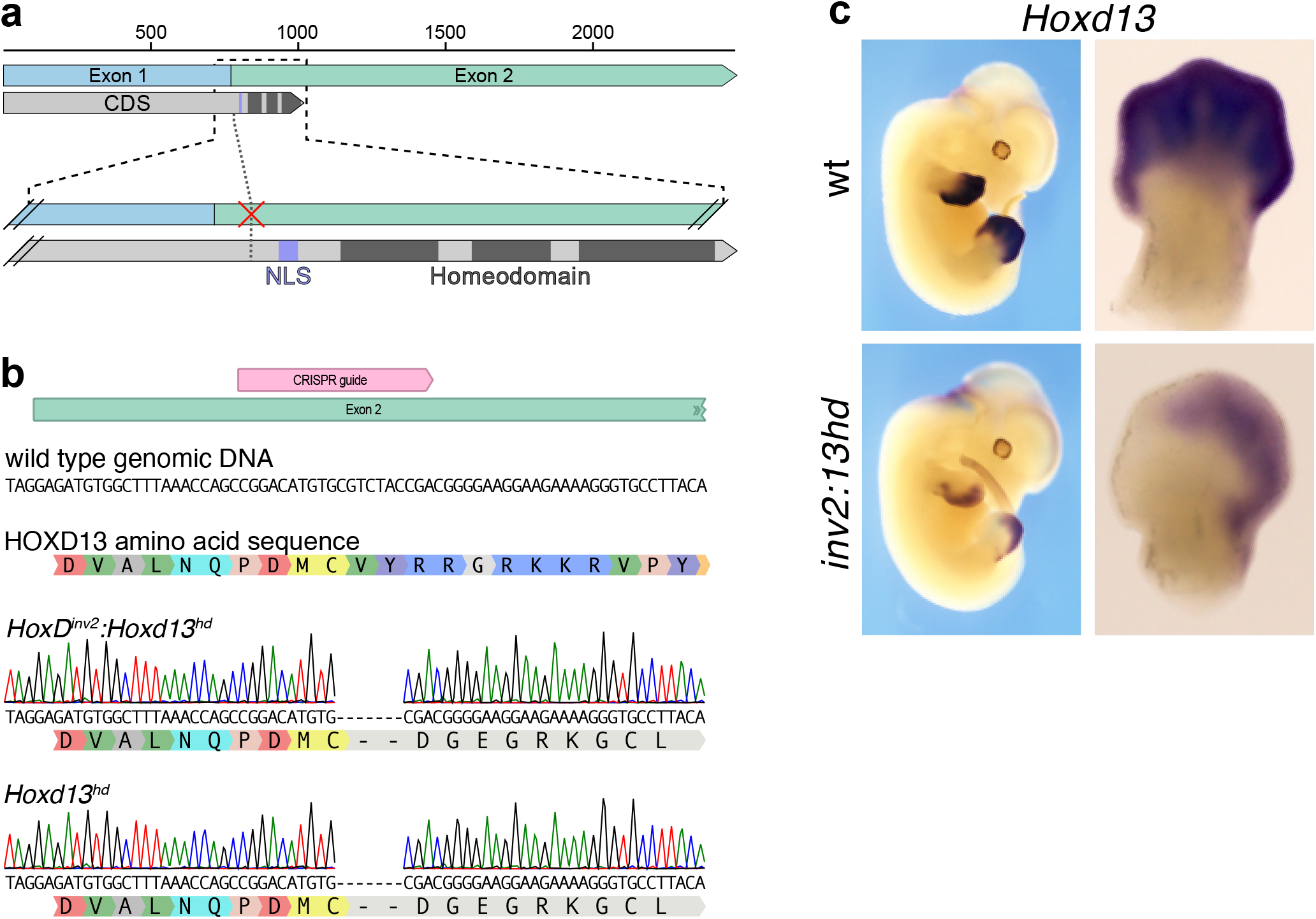
Generation and validation of the *Hoxd13^hd^* CRISPR-induced mutation. (a) Map of the *Hoxd13* spliced mRNA (top panel, blue and green) with the protein coding sequence below (grey), with the three alpha-helices of the homeodomain (dark grey). The bottom panel indicates the position of the CRISPR guide that is shown with a red X, N-terminal to the nuclear localization domain (NLS). (b) Two alleles were generated using the same CRISPR guide. The first one had a 7bp deletion in exon 2, in *cis* with the *HoxD^inν2^* allele (*HoxD^inν2^:Hoxd13^hd^*). The second allele had the same 7bp deletion, yet on the wild type chromosome (*Hoxd13^hd^*). In both cases, a frame-shift had occurred starting N-terminal to the NLS, which produced a non-functional homeodomain. (c) *In situ* hybridizations for *Hoxd13* on E12.5 *HoxD^inν2^:Hoxd13^hd^* embryos, showing that the secondary mutation did not alter location or the quantity of mRNA when compared to the initial *HoxD^inν2^* allele (see Figure 1c).

**Supplementary Figure 2.**
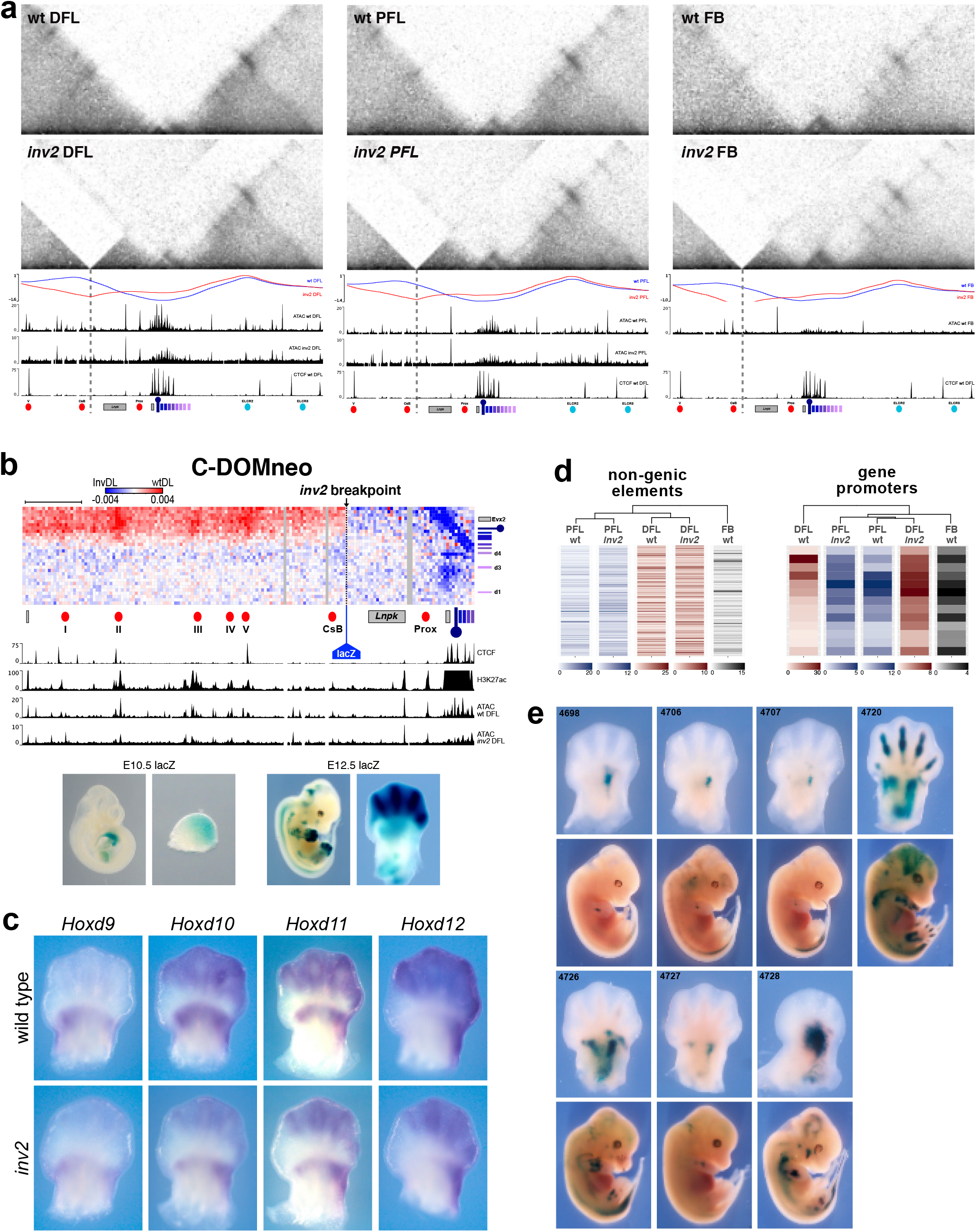
Supplementary panels for experiments in Figure 2. (a) Control and mutant conditions for Capture Hi-C for the region around the *Hoxd* gene cluster (chr2:74277600-75147000). Bin size is 5kb, color scale is log transformed. CHi-C is mapped onto wild type mm10 in the *inν2* samples. The centromeric breakpoint position is indicated by a grey dashed line. The telomeric breakpoint is not visible at this scale. The red and blue lines below the CHi-C heatmaps are TAD-separation scores. The two ATAC-Seq tracks for the corresponding tissues (only DFL and PFL) are shown along with CTCF CUT&RUN for wt DFL. (b) Subtraction of contact frequencies in CHi-C in *inν2* distal limb (DL) samples from wild type DL. These contacts are mapped on the wild type mm10 genome comparing two regions (x-axis chr2:73900700-74703000 against y-axis chr2:74638000-74783000) demonstrates the changes in contacts between the *Hoxd* gene cluster and the C-DOM created by the *inν2* allele. Red indicates greater contact frequency in the wild type distal limb and blue is greater contact in the *inν2* distal limb. The inversion creates novel regulatory environment that merges several proximal limb and distal limb enhancers. A *lacZ* sensor at the position of the inversion breakpoint responds to both types of enhancers and so the *lacZ* is detected in proximal and distal limb domains (bottom panel). (c) WISH on E12.5 forelimbs for *Hoxd* genes show very minor changes in gene expression in the limbs for these genes. *Hoxd12, Hoxd11*, and *Hoxd10* are reduced in the distal limb. In the proximal limb *Hoxd10* and *Hoxd11* are slightly reduced in the central region. On the posterior region of the proximal limb *Hoxd11* and *Hoxd12* show a very slight increase. (d) Pearson correlation hierarchical clustering on the ATAC datasets in Figure 2. The color of each bin represents the average read density in that region. In the left panel, this analysis evaluates the ATAC profiles on the non-genic portions of the *HoxD* locus. Each sample clustered most closely based on tissue of origin. The right panel uses the same clustering analysis but only evaluating the ATAC profiles on accessible *Hoxd* gene promoters. The promoters of *Hoxd* genes in the two *inν2* samples cluster closely with the wild type PFL sample indicating changes in the accessibility of their promoters in the inversion configuration. (e) All embryos collected from the PLE:lacZ transgenic experiment (Figure 2b).

**Supplementary Figure 3.**
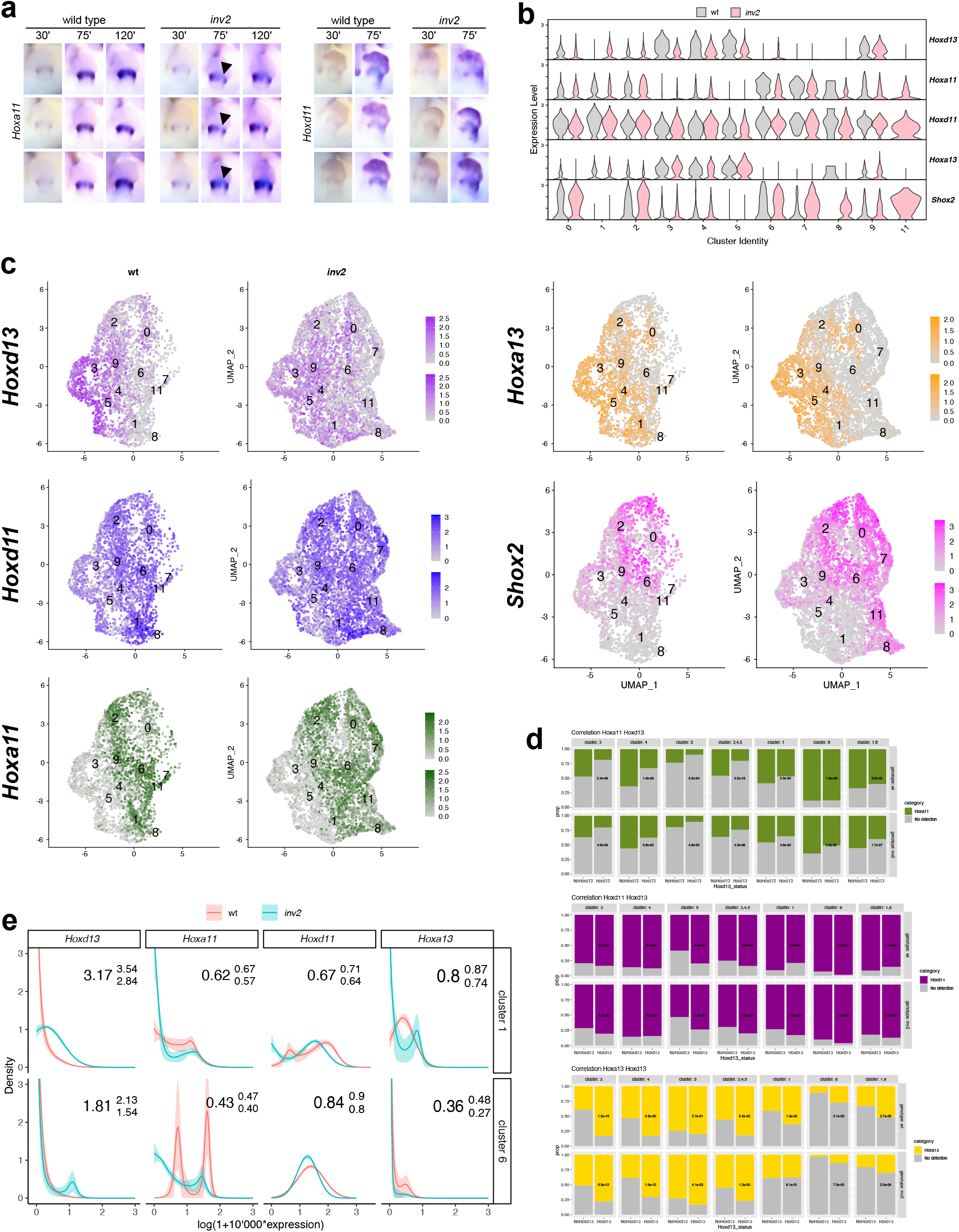
Supplementary panels for experiments in Figure 3. (a) *In situ* hybridizations for *Hoxa11* and *Hoxd11* at E12.5 through a staining time-course. In triplicate, wild type and *inν2* embryos were stained with *Hoxa11* or *Hoxd11* probes. During the development of the stain, the embryos were photographed at 30 minutes, 75 minutes, and 120 minutes (*Hoxa11* only) in order to identify any region in the proximal limb that develops less or more slowly than the wild type limbs. In the *Hoxa11* samples a small region of the posterior portion of the limb bud stained very slowly (arrowhead). *Hoxd11* showed a weak increase in staining in the same position as the loss of *Hoxa11*, which is likely to arise due to a *cis*-effect from the inversion, not due to a trans-effect of the ectopic HOXD13 protein. (b) Violin plots of the detected expression for the main clusters to determine which clusters have gained *Hoxd13* in the *inν2* samples but also express *Hoxa11* and *Hoxd11* in the wild type. *Hoxa13* was used to approximate distal limb clusters and *Shox2* for the proximal limb. (c) Expression of the genes used to determine the relevant proximal limb clusters displayed on UMAP. (d) Proportion of cells with detected UMI for *Hoxa11*, *Hoxd11*, and *Hoxa13* for each cluster and each genotype, in cells where *Hoxd13* is detected (*Hoxd13*) or not detected (No*Hoxd13*). Numbers indicate the p-value for the Fisher’s exact test. (e) Distribution of inferred true expression of *Hoxd13*, *Hoxa11*, *Hoxd11* and *Hoxa13*, for cluster 1 and cluster 6 in each genotype. The solid line indicates the mean distribution while the shaded area indicates the 68% confidence interval. In the right corners are indicated an estimation and the 68% confidence interval of the fold-change of mean expression.

**Supplementary Figure 4.**
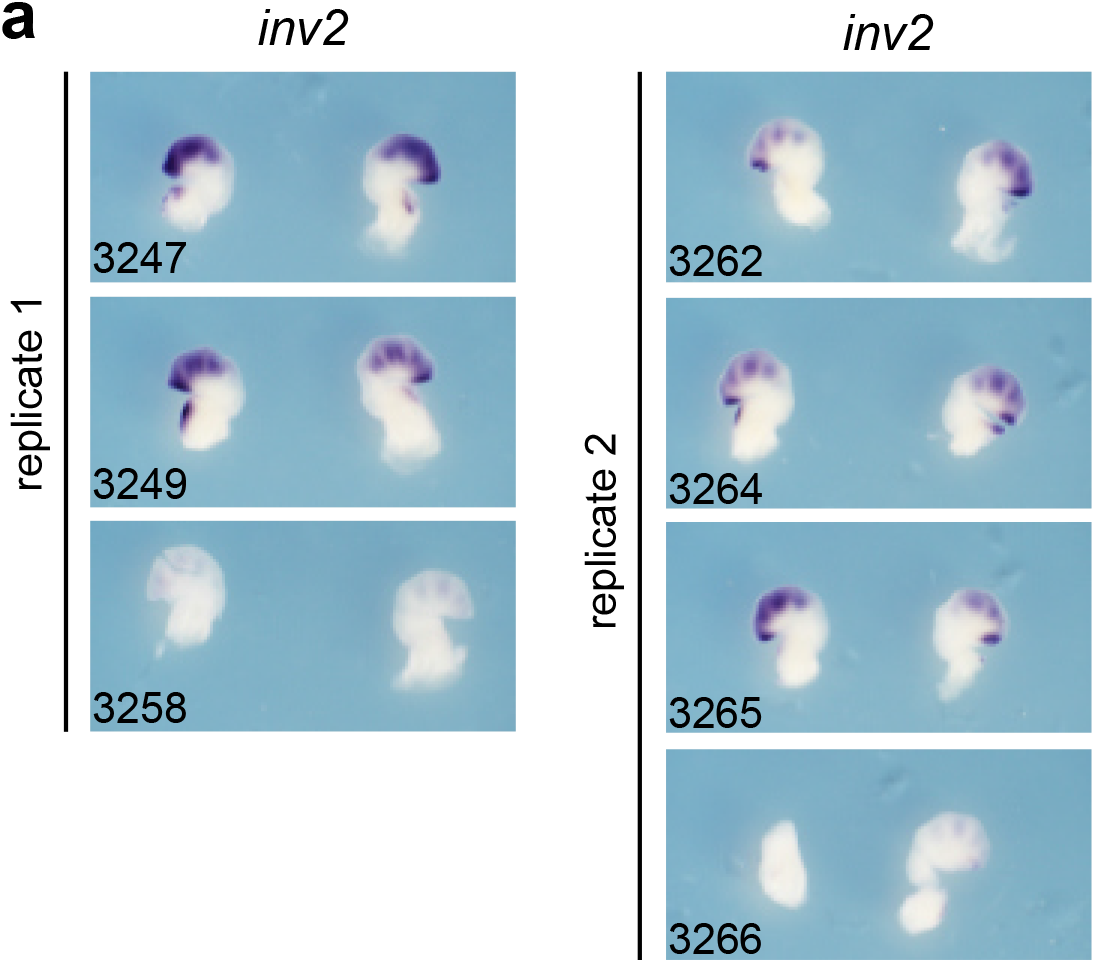
Embryos used for the CUT&RUN experiment with HOXD13. Limbs were kept after micro-dissection of the posterior proximal forelimb. They were then processed for WISH with a *Hoxd13* probe to estimate in retrospect how much distal limb contamination was present in the *inν2* replicates. The sample shaker failed during probe incubation leading to the variation in staining.

**Supplementary Figure 5.**
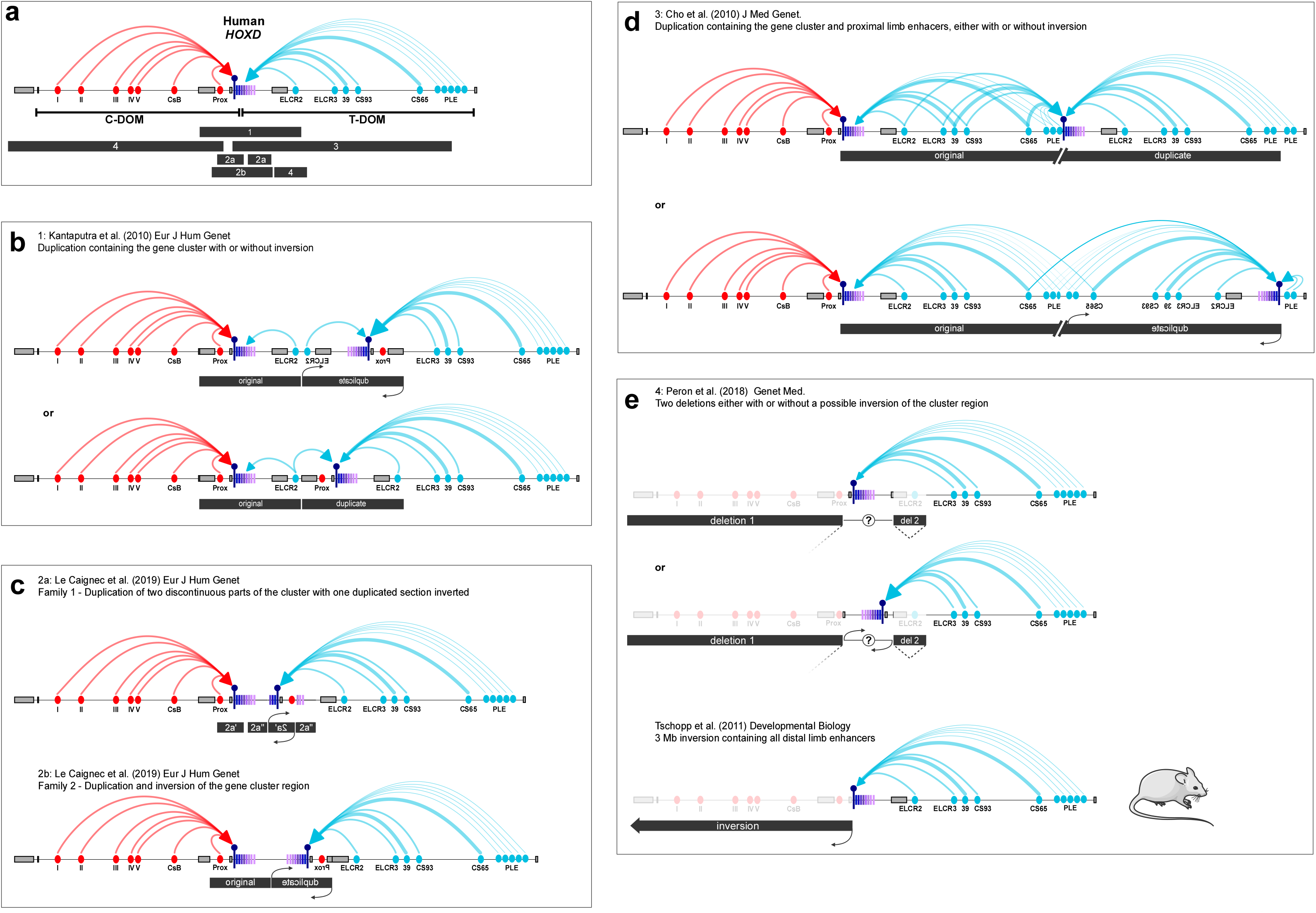
An inclusive mechanistic model for all human mesomelic dysplasia associated with 2q31. (a) Recapitulated scheme of the human regulatory C-DOM and T-DOM landscapes flanking the *HOXD* cluster, with the distal (red) and proximal (blue) regulations. The black rectangles below indicate the positions of the various chromosomal rearrangement that are detailed under panels (b-e). (b) Kantaputra et al. (2010) reported a duplication of unknown orientation (either top or bottom). In both cases, the duplicated copy of *HOXD13* (purple pin) is moved away of the distal enhancers, which continue to activate the native *HOXD13* copy, and positioned at the vicinity of the proximal enhancers, certainly leading to its ectopic expression in proximal limbs and concurrent mesomelic dysplasia. (c) In the two families reported by Le Caignec et al. (2019), the same explanatory framework can be used. In the first family (top), a double duplication with one inversion brings *HOXD13* right next to the proximal limb enhancers, while distal enhancers are still involved in the regulation of the native *HOXD13* in distal cells. In family 2 (bottom), a simpler inverted duplication has the exact same effect. (d) The condition reported by Cho et al. (2010) is slightly more complicated as a duplication of unknown orientation includes both the gene cluster and most of T-DOM i.e. the proximal regulatory landscape. In both cases, however, it is clear that *HOXD13* must fall under the control of proximal enhancers, either from both sides (top, non-inverted duplication), or from one side only (bottom, inverted duplication), leading again to a proximal gain of expression and concurrent mesomelic dysplasia. (e). The case reported by Peron et al. (2018) involves two separate deletions, either in *cis* or in *trans* (undetermined). The larger deletion (deletion 1) removes all of the C-DOM containing the distal enhancers, whereas a shorter deletion (del 2) removes a piece of the T-DOM. Should the deletions be in *cis*, the orientation of the cluster located in between could be either way, as a result of the two deletions (Kragesteen et al., 2018), thus leading to two potential configurations (top and middle). The first configuration (top) is very similar to an engineered allele produced in mice (bottom) whereby all distal enhancers were separated from the *HoxD* cluster through a large inversion (Tschopp and Duboule, 2011). Constitutive contacts between *Hoxd13* and some of these enhancers were released, thus allowing *Hoxd13* to interact with proximal enhancers and to be expressed in proximal cells leading to a weak mesomelic dysplasia (Montavon et al., 2012; Tschopp and Duboule, 2011). The exact same phenomenon would likely occur in configuration 1 of Peron et al. (2018). Should the deletions be in *cis* and the cluster inverted (middle), the latter effect would be strengthened by an increased proximity between *HOXDl3* and proximal enhancers.

---

**Supplementary Table 1S**. The supplementary table contains: the genotyping primer sequences for the *HoxD^inν2^* and *Hoxd13^hd^* alleles, the CRISPR guide sequence used to generate the *Hoxd13^hd^* allele, the genomic coordinates for the enhancer sequences used in the PLE:lacZ transgene assay, and the DNA sequence for the PLE:lacZ enhancer-reporter construct.

## METHODS

### Animal work

All experiments were performed in agreement with the Swiss Law on Animal Protection (LPA) under license numbers GE 81/14 and VD2306.2 (to D. Duboule).

### Generation of the *HoxD^inν2^* allele

The *HoxD^inν2^* allele was generated by STRING (Spitz et al., 2005) using a cross between mice carrying the del65 allele (Andrey et al., 2013) and mice carrying a loxP site inserted at chr2:74477755 (mm10) by a Sleeping Beauty transposon system (Ruf et al., 2011). F0 mice carrying these two loxP sites in-*cis* were then crossed with the *Hprt*-Cre (Tang et al., 2002) and F1 animals from this cross were screened for the presence of the inversion between the coordinates chr2:74477755-75438813 (mm10) with genotyping primers included in Supplementary Table 1. The regions of the inversion breakpoints were confirmed by Sanger sequencing.

### Generation of the secondary *Hoxd13^hd^* mutation

We used a single CRISPR guide sequence that was used previously for the mutation of *Hoxd13* (Supplementary Table 1)(Darbellay et al., 2019). The guide sequence was transcribed *in vitro* with NEB HiScribe T7 (NEB E2040S). The guide and TrueCut Cas9 v2 protein (ThermoFisher A36497) were electroporated with a NEPA21 (NEPA GENE Co. Ltd., Chiba, Japan) into fertilized mouse embryos carrying the *HoxD^inν2^* allele as previously reported (Kaneko et al., 2014). Founders were screened by PCR for the presence of the *inν2* allele and mutation to the *Hoxd13* second exon. Each founder was crossed with an allele containing a deletion of the *Hoxd* gene cluster to determine which founders contain the *inν2* and *Hoxd13^hd^* mutations in-*cis* and we also screened for alleles containing the *Hoxd13^hd^* mutation on the wild type chromosome to use as controls. A founder stock was identified for both alleles containing the same DNA mutation. Micro-CT scans were performed on littermates at five weeks after birth at the University of Geneva CMU and analyzed with Horos 3.3.6. Box plots for length measurements and t-test were produced in DataGraph 4.6.1.

### Histology, *In situ* hybridizations, and *lacZ* stains

Embryos were collected at the indicated stages and processed as previously reported (Woltering et al., 2009). Embryos were treated with Proteinase K (EuroBio GEXPRK01-15): E10.5 and E11.5 at 1:2000 for 7 min, E12.5 at 1:1000 for 10 min, E13.5 at 1:1000 for 14 min, E14.5 at 1:1000 for 40 min. For P3 skeletons, animals were collected at post-natal day 3. Alcian Blue and Alizarin Red stains were performed as previously reported (Rigueur and Lyons, 2014). *lacZ* stains were performed as previously reported (Yakushiji-Kaminatsui et al., 2018).

### ATAC-Seq

Embryos were collected at E12.5 and placed in PBS on ice. Yolk sacs were collected, digested in buffer (10mM EDTA pH8.0 and 0.1mM NaOH) at 95° for 10 min with shaking at 900rpm. DNA from these samples was screened by genotyping with Z-Taq (Takara R006B). Embryos identified as homozygous for wild type or *HoxD^inν2^* alleles were processed individually for ATAC-Seq. The proximal forelimbs of embryos were dissected (see Figure 3) and placed into PBS with 10% FCS and 8ul collagenase at 50mg/ml (Sigma C9697) at 37° for approximately 5min. Cells were counted and 50,000 cells were isolated for processing with the Nextera Tn5 enzyme (Illumina FC-131-1096) as previously described (Buenrostro et al., 2013). Tn5 treated DNA was amplified with Nextera Library primers using NEBNext library amplification master mix (NEB M0541) and sequenced on an Illumina NextSeq. Sequenced DNA fragments were processed as previously reported (Amândio et al., 2020) with a minor modification: the BAM file was converted to BED prior to peak calling with bedtools version 2.27 (Quinlan 2010). Peak calling was done using MACS2 (v2.1.1.20160309) callpeak (--no-model --shift −100 -- extsize 200 --call-summits --keep-dup all). For hierarchical clustering and heatmap analysis, two sets of bed files were generated. First, the peak regions were collected from the three wild-type ATAC datasets, then merged with bedtools (version 2.27.1, citation) to remove redundant elements. The merged peaks in the region chr2:73950000-75655000 excluding the *Lnpk* and *Mtx2* gene bodies as well as the region from *Evx2* to *Hoxd1* constitute the first set composed of non-genic elements. The second one contains the promoters (−1kb, +100bp from TSS) that overlapped with at least one peak in a wild-type ATAC dataset in the region from *Evx2* to *Hoxd1*. Heatmaps were generated with plotHeatmap from deepTools version 3.5 (Ramírez et al., 2016). Clustering was performed with R (www.r-project.org) on matrices generated by multiBigWigSummary (deepTools version 3.5).

## CUT&RUN

Posterior proximal forelimb cells were isolated and genotyped in the same manner as samples for the ATAC-Seq (above). Pools of cells from individual embryos (see Figure 4S) were processed according to the CUT&RUN protocol (Skene et al., 2018) using a final concentration of 0.02% digitonin (Apollo APOBID3301). Cells were incubated with 0.5ug of anti-HOXD13 antibody (Abcam ab19866), or anti-CTCF (Active Motif 61311) at 4°C. The pA-MNase was kindly provided by the Henikoff lab (Batch #6) and added at 0.5ul/100ul Digitonin Wash Buffer. Cells were digested in Low Calcium Buffer and released for 30 minutes at 37°C. Sequencing libraries were prepared with KAPA HyperPrep reagents (07962347001) with 2.5ul of adaptors at 0.3uM and ligated for 1 hour at 20°C. The DNA was amplified for fourteen cycles. Post-amplified DNA was cleaned and size selected using 1:1 ratio of DNA:Ampure SPRI beads (A63881) followed by an additional 1:1 wash and size selection with HXB. HXB is equal parts 40% PEG8000 (Fisher FIBBP233) and 5M NaCl. Sequenced DNA fragments were processed as previously reported (Amândio et al., 2020) with slight modifications: PCR duplicates were removed with picard before the BAM to BED conversion and in MACS2 using the option --keep-dup all instead of --keep-dup 1. Motif enrichment was performed on individual samples with HOMER version 4.10 (Heinz et al., 2010) using default conditions on peaks identified as significant from MACS2 in the second replicate. All samples were mapped to wild type mm10. The E11.5 whole forelimb HOXA11 ChIP-Seq datasets (Desanlis et al., 2020)(SRR8290670 and SRR8290672) were down-sampled to 25mio reads with seqtk version 1.3 (https://github.com/lh3/seqtk/) using a RNG seed of 4 and then processed following a previously reported workflow (Beccari et al., 2020). Differential binding analysis was performed with DiffBind 2.14.0 (Ross-Innes et al., 2012) on replicate sample peak sets identified by MACS2 for HOXD13 in wild type distal forelimb and HOXA11 in wild type forelimb, with default conditions using DESeq2 1.24.0. Hierarchical clustering analysis was performed with deepTools plotCorrelation, and the heatmap was generated with deepTools plotHeatmap.

### Capture Hi-C

Samples used in the Capture Hi-C were identified by PCR screening embryos at E12.5 as described above. Collagenase treated samples were cross-linked with 1% formaldehyde (ThermoFisher 28908) for 10 minutes at room temperature and stored at −80° until further processing as previously described (Yakushiji-Kaminatsui et al., 2018). The SureSelectXT RNA probe design used for capturing DNA was done using the SureDesign online tool by Agilent. Probes cover the region chr2:72240000-76840000 (mm9) producing 2x coverage, with moderately stringent masking and balanced boosting. Sequenced DNA fragments were processed as previously reported (Yakushiji-Kaminatsui et al. 2018) but the mapping was performed on mm10 and the reads in chr2:72402000-7700000 were selected. The mutant *inν2* genome was characterized from Sanger sequencing data around the inversion breakpoints. A custom R (www.r-project.org) script based on the SeqinR package (Charif and Lobry, 2007) allowed the construction of a FASTA file for the inverted chromosome 2 from the wild-type sequence and the exact position and sequence of breakpoints. The sequence of the mutant chromosome 2 (available at 10.5281/zenodo.4456654) was then compiled with other wild-type chromosomes to form the mutant *inν2* genome. For samples that were mapped to the *inν2* mutant genome, the same workflow as described above was used. Heatmaps in Figure 2 were plotted with pyGenomeTracks 3.3 (Lopez-Delisle et al., 2020; Ramírez et al., 2018) and subtraction heatmaps in Figure 2 and 2S were plotted with a custom tool available at https://github.com/lldelisle/scriptsForBo1tEtA12021. The TAD separation scores in Figure 2S were computed with HiCExplorer hicFindTADs version 3.5.1.

### Enhancer transgenesis assay

The enhancer regions used in the transgenesis assay (Figure 2) were identified based on the presence of H3K27Ac histone modification and ATAC peaks in wild type proximal forelimbs, and also on the absence of CTCF. DNA sequences from these regions (Supplementary Table 1) were collected and assembled *in silico* to produce the PLE TgN sequence with *KpnI* and *ApaI* restriction sites flanking the enhancer sequences. This 3.5 kb DNA sequence was synthesized by TWIST Bioscience (San Francisco, CA). The PLE sequence was restriction digested and ligated into the *pSK-lacZ* reporter construct. The 7.1 kb fragment containing the PLE:*lacZ* construction was excised from the vector backbone with the *KpnI* and *SacII* restriction enzymes, purified by agarose gel and column purification (Qiagen 28704). Pro-nuclear injections were performed by the University of Geneva CMU. Embryos were collected at approximately E12.5 and stained for *lacZ*.

### Single cell RNA-seq

Embryos were collected and stored in 1X PBS treated with DEPC and held on ice while genotyping was performed. Embryos with the desired genotype were selected and the posterior portion of each of the forelimbs was isolated for each replicate. The cells were digested in collagenase and stored in 1X PBS containing 10% FCS and 0.2mM EDTA to prevent cellular aggregation. The cells samples were transferred to the EPFL Gene Expression Core Facility for preparation into 10X GEMs and reverse transcription according to the 10X Chromium 3.1 protocol. Sequencing reads were processed with Cell Ranger 3.1.0 for demultiplexing, barcode processing, and 3’ gene counting using a modified gene annotation file (10.5281/zenodo.4456702). Clustering analysis was performed with Seurat 3.2.3 with R 3.6.3. All commands used are available (https://github.com/lldelisle/scriptsForBo1tEtA12021). In order to evaluate the inferred true distribution of expression, not the distribution of detected expression, we used a new method based on Monte-Carlo Markov Chain (Lopez-Delisle and Delisle manuscript in preparation). Briefly, this method assumes that the original distribution of expression (probability density function: pdf) can be approximated by a number of Gaussians provided by the user. We evaluate the likelihood of the Gaussian parameters by using the raw expression and the total number of UMI observed for each cell assuming they follow a Poisson distribution. Then, we do a Markov chain Monte Carlo (MCMC) to obtain an interval confidence for the pdf. In order to compare models obtained with different number of Gaussians, we use the model with the highest value of evidence. From these pdf, we can also get mean expression for each sample of the MCMC and thus evaluate a confidence interval of the fold-change between two conditions. This has been used in Supplementary Figure 3e where median and 68% confidence interval is given. This method also extends for 2 genes. Instead of having a pdf in 1 dimension for 1 gene, the pdf is now in 2 dimensions (2d) corresponding to the two genes. The pdf is approximated by a given number of 2d Gaussians. In addition to an estimation of the pdf in 2d, we also get a confidence interval of the correlation value and we can evaluate a confidence interval on the one-sided p-value. In this analysis, we used MCMC of 1 million samples, the axis (x or y) was divided in 100 bins for Supplementary Figure 3e and 50 bins for Figure 3f, all combinations of numbers of Gaussian between 1 and 4 (1 and 5 for Supplementary Figure 3e) were tested and only the best model was kept. The results were processed in R and plotted with ggplot2 (Wickham, 2016).

---

**Supplementary Table 1.**
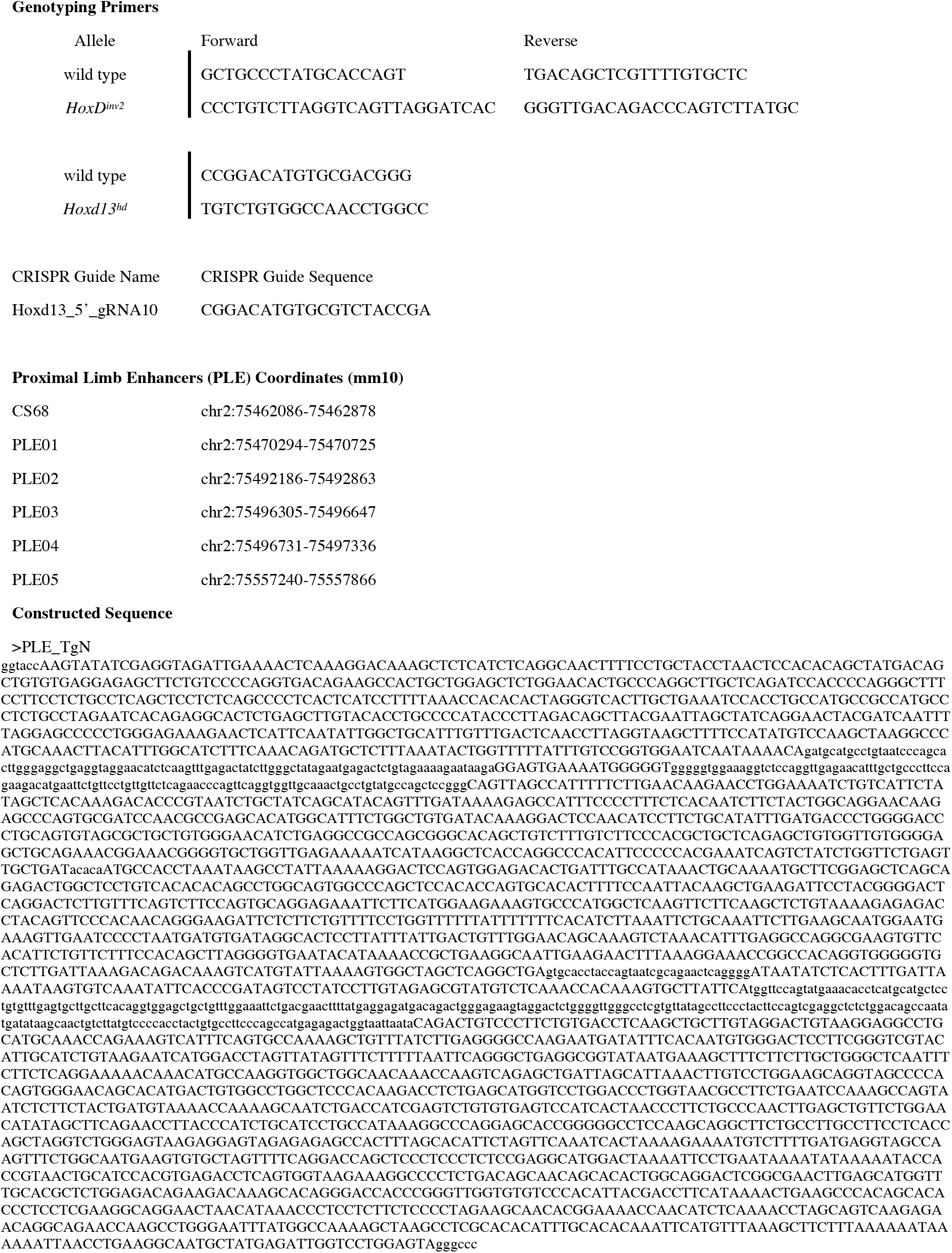
List of genotyping primers, CRISPR guide sequences, and enhancer elements.

## Supplementary Video 1

Micro-CT scans of adult left forearm skeletons. All skeletons are homozygous for the indicated genotype.

https://drive.switch.ch/index.php/s/UJpBrsRbI1KY6qo

